# Mitochondrial protein import stress causes progressive neurodegeneration opposed by PERK - eIF2α signalling

**DOI:** 10.64898/2026.04.10.717732

**Authors:** Johannes Ebding, Marlene Barth, Lena Maria Lion, Adrian Gackstatter, Sarah Link, Marcello Pirritano, Gilles Gasparoni, Martin Simon, Johannes M. Herrmann, Jan Pielage

## Abstract

Mitochondrial dysfunction and impairments of the mitochondrial protein import system are often linked to neurodegenerative disease, but whether import stress per se causes neurodegeneration has not been tested. Here, we adapted the yeast clogger system to Drosophila motoneurons to block TOM-TIM23-mediated import with temporal control. Sustained import stress converts somatic mitochondria into donut-shaped structures, depletes functional mitochondria from synaptic terminals, and causes progressive neurodegeneration with impaired neurotransmitter release and locomotor dysfunction. This neurodegeneration is mechanistically distinct from mitochondrial absence, as miro mutant neurons that completely lack presynaptic mitochondria do not degenerate. Import-stressed motoneurons activate multiple protective programmes, including chaperone remodelling, metabolic repression, and translational control through the eIF2α kinase PERK. Both pharmacological PERK inhibition and reversal of translational attenuation via ISRIB accelerate neurodegeneration, whereas PERK overexpression alone is sufficient to cause it, defining a protective range of eIF2α-dependent translational control. The observation that PERK inhibition is protective in protein misfolding models but detrimental during import stress shows that the nature of mitochondrial dysfunction determines the molecular consequence of translational control in neurodegeneration.

## Introduction

Mitochondrial biogenesis depends on the post-translational import of more than 1,000 nuclear-encoded proteins through dedicated translocases (Pfanner *et al*, 2025). Genetic evidence links this process to neurological disease. Germline mutations in the translocase subunits *TIMM8A* and *TOMM70* cause deafness-dystonia syndrome and severe neurological impairment, respectively (Koehler *et al*, 1999; Dutta *et al*, 2020; Wei *et al*, 2020), indicating that the nervous system is particularly sensitive to reduced mitochondrial import capacity. Several proteins associated with common neurodegenerative diseases physically obstruct import channels. Amyloid precursor protein accumulates in the TOM-TIM23 supercomplex in Alzheimer’s disease brain tissue, and Aβ peptides impair preprotein import competence through coaggregation with precursor proteins (Devi *et al*, 2006; Cenini *et al*, 2016). α-Synuclein associates with TOM20 and inhibits import in Parkinson’s disease models and post-mortem tissue (Di Maio *et al*, 2016), and mutant huntingtin binds TIM23 and inhibits mitochondrial protein uptake (Yano *et al*, 2014; Yablonska *et al*, 2019). The most direct evidence comes from pathogenic mutations in the adenine nucleotide translocase ANT1, which cause the protein to arrest within the TOM-TIM22 import pathway, producing neurodegeneration and dominant myopathy in mice (Coyne *et al*, 2023). In each of these diseases, however, import stress cannot be separated from protein aggregation, gain-of-function toxicity, or loss of the disease protein’s normal cellular role. Whether mitochondrial protein import stress, as the sole cellular insult, is sufficient to cause neurodegeneration, and how neurons respond to this insult in the absence of these additional mechanisms, remain open questions.

How cells respond to sustained import stress has been systematically dissected in yeast, revealing a hierarchy of defences that escalate with the severity and duration of the blockade. When import is acutely impaired, cells activate a transcriptional programme through the heat shock factor Hsf1, inducing the major cytosolic chaperone Hsp70 to stabilise accumulating precursor proteins and prevent their misfolding (Boos *et al*, 2019). In parallel, the transcription factor Rpn4 drives upregulation of the ubiquitin-proteasome system to degrade precursors that cannot be refolded (Wrobel *et al*, 2015; Wang & Chen, 2015; Boos *et al*, 2019), and a dedicated extraction machinery removes stalled precursors directly from clogged translocons (Mårtensson *et al*, 2019; Weidberg & Amon, 2018). As import stress persists and proteasomal capacity is exceeded, chaperone-controlled cytosolic granules termed MitoStores sequester excess precursors in a reversible, non-toxic form (Krämer *et al*, 2023). If these protective mechanisms fail, non-imported precursors, which are metastable and aggregation-prone, seed the co-aggregation of other cytosolic proteins, including disease-associated species such as α-synuclein and amyloid β (Nowicka *et al*, 2021). In mammalian cells, mitochondrial stress is relayed to the cytosol through a dedicated pathway in which the protease OMA1, its substrate DELE1, and the eIF2α kinase HRI activate the integrated stress response (Guo *et al*, 2020; Fessler *et al*, 2020; Haakonsen *et al*, 2024). How neurons respond to sustained import blockade, however, is largely unknown. Whether the yeast-defined quality control hierarchy operates in neurons, and whether it protects the synapse or contributes to its degeneration, has not been tested.

Neurons face qualitatively different constraints in managing import stress. In dividing cells, precursor proteins that accumulate in the cytosol during import blockade are diluted with each cell division; post-mitotic neurons lack this mechanism and must clear or tolerate the entire precursor burden cell-autonomously. Neuronal polarity imposes an additional constraint: mitochondria must be transported from the soma, where the majority of protein synthesis occurs, to synaptic terminals that lie at the end of axons orders of magnitude longer than the cell body (Guo *et al*, 2005). Any impairment that alters mitochondrial morphology or fitness in the soma therefore has the potential to restrict mitochondrial delivery to the compartment that depends on it most. The synaptic terminal, in turn, requires continuous local ATP production to sustain vesicle cycling (Rangaraju *et al*, 2014), and continuous delivery of newly synthesised maintenance proteins to preserve its structural integrity. This dependency creates a tradeoff that does not arise in yeast: a cellular response that attenuates translation to limit precursor production simultaneously threatens the supply of proteins on which synaptic maintenance depends. Whether neurons resolve this tradeoff, and how, has not been addressed, in part because no experimental system has generated sustained import blockade in neurons in vivo while isolating import stress as the sole cellular insult.

Here, we adapted the yeast clogger system (Boos *et al*, 2019) to *Drosophila* motoneurons. The larval neuromuscular junction provides genetic access to all neuronal compartments, from soma through axon to synaptic terminal, with established assays for synaptic integrity and degeneration at single-synapse resolution, electrophysiological function, and locomotor behaviour (Pielage *et al*, 2005; Mushtaq *et al*, 2022). A mitochondrial-targeted DHFR-based construct stalls in the TOM-TIM23 import channel and competitively blocks import of endogenous precursor proteins (Boos *et al*, 2019). Tissue-specific and temporal control via the UAS-GAL4-based TARGET system (McGuire *et al*, 2003) allows staging of the degenerative response from onset through progression. We show that sustained import stress causes progressive synaptic degeneration that is mechanistically distinct from both mitochondrial absence and respiratory chain deficiency. The transcriptomic response shares core features with the yeast programme but additionally activates the integrated stress response. Pharmacological experiments demonstrate that the ISR kinase PERK opposes synaptic degeneration during import stress through eIF2α-mediated translational control. These data identify mitochondrial protein import stress as a sufficient cause of neurodegeneration in vivo and establish that the protective or detrimental outcome of translational control during neurodegeneration is determined by the nature of the upstream mitochondrial stress.

## Results

### Mitochondrial clogger constructs target mitochondria in *Drosophila* motoneurons

To selectively block mitochondrial protein import in *Drosophila* motoneurons, we adapted the clogger system established in yeast (Boos *et al*, 2019). In clogger constructs, the tightly folded mouse dihydrofolate reductase (DHFR) domain is fused to a mitochondrial targeting sequence (MTS). Because the folded DHFR domain cannot pass through the TOM translocation pore, it stalls within the channel and competitively blocks import of endogenous precursor proteins. We generated three GFP-tagged UAS constructs by phiC31-mediated transgenesis (Bischof *et al*, 2007) at the attP40 landing site to ensure equivalent expression levels (Figure 1A): cyt^DHFR^, a cytosolic control that lacks any targeting sequence; mito^DHFR^, which carries the MTS derived from cytochrome b2 of budding yeast fused to DHFR, directing the protein toward the mitochondrial matrix; and mito-IMM^DHFR^, which carries the same MTS but additionally retains the transmembrane domain of cytochrome b2, serving as a stop-transfer signal that arrests the protein at the inner mitochondrial membrane. In yeast, both constructs occupy the TOM-TIM23 import channel and compete with endogenous precursors, but the transmembrane anchor in mito-IMM^DHFR^ prevents its release and causes sustained channel blockade, whereas mito^DHFR^ produces a milder import impairment (Boos *et al*, 2019).

**Figure 1:**
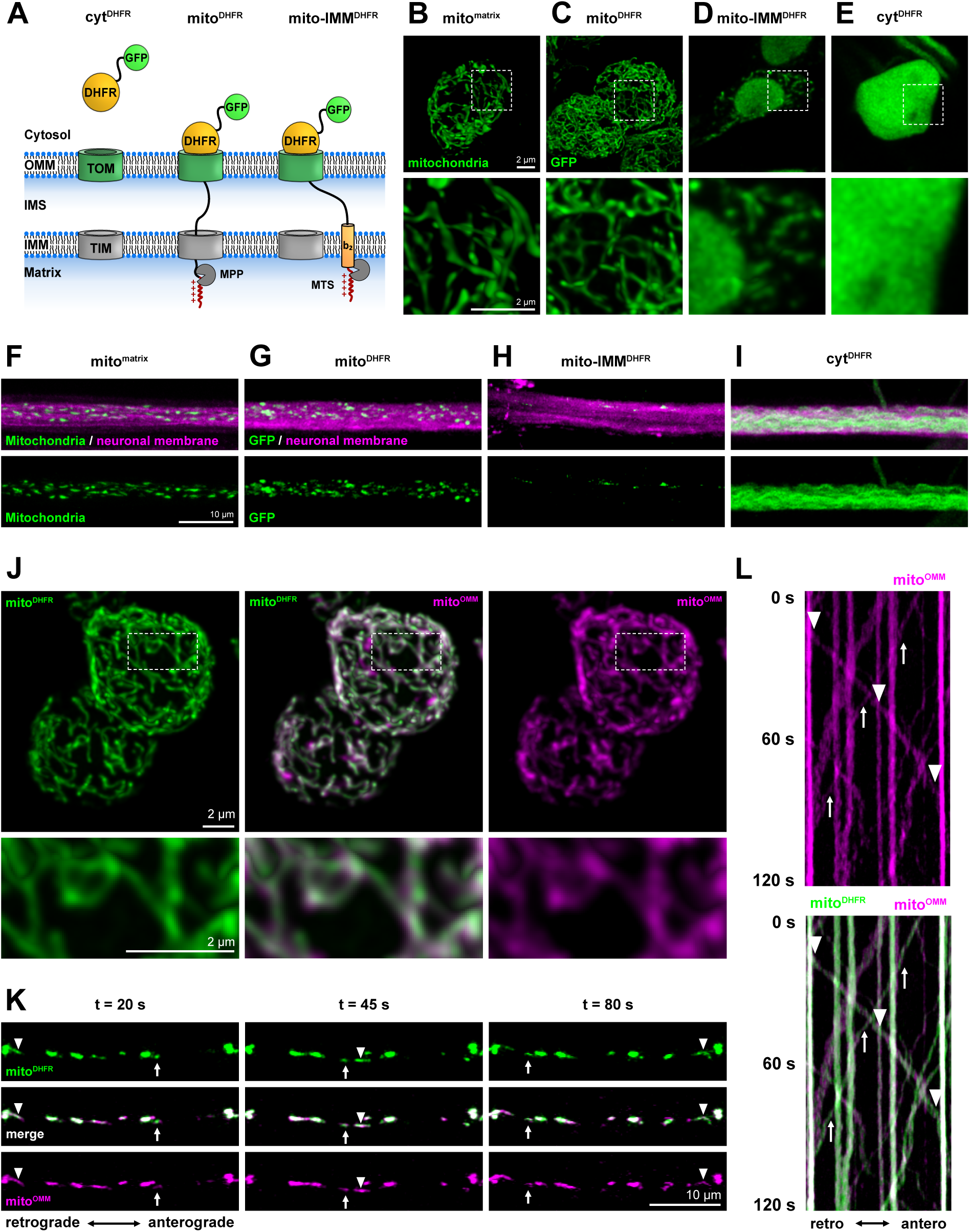
Design and localisation of mitochondrial clogger constructs in *Drosophila* motoneurons. **(A)** Schematic of clogger construct design. All constructs contain a GFP tag and were integrated at the attP40 landing site by phiC31-mediated transgenesis. cyt^DHFR^ lacks a mitochondrial targeting sequence (MTS) and remains cytosolic. mito^DHFR^ carries the MTS of cytochrome b2 fused to the dihydrofolate reductase (DHFR) domain, directing it toward the mitochondrial matrix; the DHFR domain cannot pass through the TOM translocation pore but may dislocate after cleavage of the MTS by the mitochondrial processing peptidase (MPP). mito-IMM^DHFR^ carries the same MTS but retains the transmembrane domain (TMD) of cytochrome b2, which serves as a stop-transfer signal anchoring the protein at the inner mitochondrial membrane and preventing release from the TOM-TIM23 channel. **(B)**–**(E)** Live imaging of GFP-tagged constructs in motoneuron somata (5053-Gal4). **(B)** The mitochondrial matrix marker mito^matrix^ labels an interconnected tubular mitochondrial network. **(C)** mito^DHFR^ shows an identical network distribution. **(D)** mito-IMM^DHFR^ partially labels the mitochondrial network with additional nuclear accumulation. **(E)** cyt^DHFR^ distributes uniformly throughout the cytoplasm with partial nuclear accumulation. Scale bars in (B) apply to (B)–(E): 2 µm. **(F)**–**(I)** Distribution of constructs in motoneuron axons (OK371-Gal4). **(F)** mito^matrix^ labels discrete individual mitochondria evenly distributed throughout nerve bundles. **(G)** mito^DHFR^ shows an identical axonal distribution. **(H)** mito-IMM^DHFR^ signal is sparse and detectable only at elevated imaging sensitivity. **(I)** cyt^DHFR^ labels all axons uniformly. Scale bars in (F) apply to (F)–(I): 10 µm. **(J)**–**(L)** Co-expression of mito^DHFR^ (green) and the import-independent outer mitochondrial membrane marker mito^OMM^ (magenta) in motoneurons (5053-Gal4). **(J)** Both markers co-localise within the mitochondrial network in the soma. Scale bar: 2 µm. **(K)** Snapshots from live imaging of an individual motoneuron axon demonstrating co-transport of both markers in anterograde and retrograde directions. Scale bar: 10 µm. **(L)** Kymograph of the recording shown in (K); total duration 120 s.

To assess construct localisation, we compared each to the established mitochondrial matrix marker mito^matrix^, an HA-GFP fusion protein targeted to the mitochondrial matrix via an MTS (Pilling *et al*, 2006). Motoneuron-specific expression (5053-Gal4) revealed that mito^matrix^ labelled an interconnected tubular mitochondrial network within the cell soma (Figure 1B). mito^DHFR^ showed an almost identical network distribution, consistent with efficient mitochondrial targeting (Figure 1C). mito-IMM^DHFR^ displayed only partial mitochondrial network labelling with additional nuclear accumulation (Figure 1D). cyt^DHFR^ distributed uniformly throughout the cytoplasm and also partially accumulated in the nucleus (Figure 1E). In mammalian cells, DHFR undergoes SUMO-dependent nuclear import as part of the de novo thymidylate biosynthesis pathway (Woeller *et al*, 2007; Anderson *et al*, 2012), and the nuclear signal in constructs not efficiently sequestered by mitochondrial import is consistent with this dual localisation. In motoneuron axons (OK371-Gal4), the somatic mitochondrial network resolved into discrete individual mitochondria evenly distributed throughout nerve bundles, as labelled by mito^matrix^ (Figure 1F). mito^DHFR^ showed an identical axonal distribution (Figure 1G). In contrast, mito-IMM^DHFR^ signal was sparse in axons and detectable only at elevated imaging sensitivity (Figure 1H), while cyt^DHFR^ labelled all axons uniformly (Figure 1I). These data indicate that both mito^DHFR^ and mito-IMM^DHFR^ target mitochondria, but import-stressed mitochondria are less efficiently delivered to the axonal compartment.

To directly confirm mitochondrial targeting, we co-expressed mito^DHFR^ with mito^OMM^ (Vagnoni & Bullock, 2016), a TOM/TIM-independent outer mitochondrial membrane marker consisting of mCherry fused to the transmembrane domain of the mitochondrial GTPase Miro. In the cell soma, mito^DHFR^ and mito^OMM^ co-localised within the mitochondrial network (Figure 1J). Live imaging of individual motoneuron axons demonstrated co-transport of both markers in anterograde and retrograde directions (Figure 1K), as confirmed by kymograph analysis (Figure 1L). These data demonstrate that mito^DHFR^ is efficiently targeted to mitochondria in *Drosophila* motoneurons.

### Mitochondrial protein import block leads to a remodelling of mitochondrial structures

Since mito-IMM^DHFR^ showed only partial mitochondrial network labelling in the soma (Figure 1D), we next investigated its impact on mitochondrial protein import and morphology. To distinguish import-dependent from import-independent effects, we employed two complementary markers (Figure 2A): mito^matrix^, which requires TOM/TIM-mediated translocation and thus reflects mitochondrial import capacity; and mito^OMM^, which labels the entire mitochondrial population regardless of import status. In control motoneurons, both markers labelled the mitochondrial network in an almost identical pattern (Figure 2A). Co-expression of mito^DHFR^ with mito^matrix^ revealed a mitochondrial network indistinguishable from cyt^DHFR^ co-expression (Figure 2B, C). Three-dimensional quantification confirmed no significant differences in mitochondrial number, volume, or surface area per soma (Figure 2E-G), and individual mitochondrial morphology, volume, and mito^matrix^ signal intensity remained unchanged (Figure 2H-J). mito^DHFR^ therefore does not detectably perturb mitochondrial protein import or morphology in the soma. In contrast, co-expression of mito-IMM^DHFR^ significantly reduced the volume and surface area of mito^matrix^-labelled structures while increasing the number of detectable individual signals (Figure 2D-G). Analysis of individual mitochondria revealed a significant increase in sphericity alongside a drastic reduction in both volume and signal intensity (Figure 2H-J). These data indicate that mito-IMM^DHFR^ blocks TOM-mediated import and thereby prevents efficient translocation of the mito^matrix^ protein into the mitochondrial matrix.

**Figure 2:**
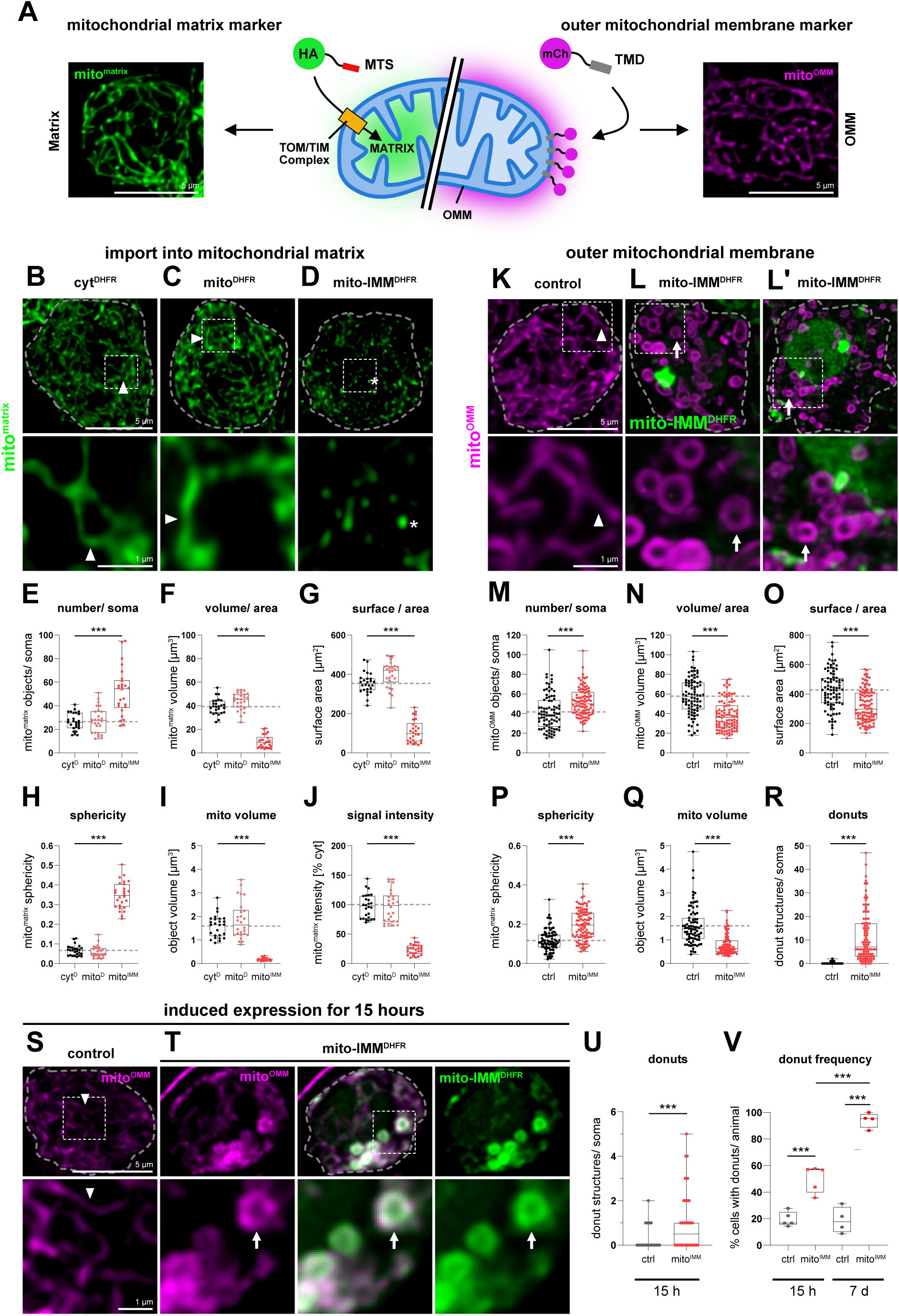
mito-IMM^DHFR^ blocks mitochondrial protein import and induces mitochondrial morphological changes. **(A)** Schematic and live imaging of the two complementary mitochondrial markers. mito^matrix^ carries a mitochondrial targeting sequence fused to HA-GFP and requires TOM/TIM-mediated translocation, reporting import capacity. mito^OMM^ carries mCherry fused to the transmembrane domain of Miro and inserts into the outer mitochondrial membrane independently of protein import, labelling the entire mitochondrial population. Both markers label the mitochondrial network in control motoneuron somata. Scale bar: 5 µm. **(B)**–**(D)** HA-staining of mito^matrix^ co-expressed with cyt^DHFR^ **(B)**, mito^DHFR^ **(C)**, or mito-IMM^DHFR^ **(D)** in motoneuron somata (OK371-Gal4). cyt^DHFR^ and mito^DHFR^ co-expression produces intact mitochondrial networks indistinguishable from controls. mito-IMM^DHFR^ co-expression disrupts the mito^matrix^ pattern, yielding fragmented, punctate signals with reduced intensity. Scale bars: 5 µm (overviews), 1 µm (insets). **(E)**–**(G)** Quantification of mitochondrial parameters per soma: number of mitochondria **(E)**, total mitochondrial volume **(F)**, and total surface area **(G)**. **(H)**–**(J)** Quantification of individual mitochondria: sphericity **(H)**, volume **(I)**, and mito^matrix^ signal intensity **(J)**. mito-IMM^DHFR^ expression significantly alters all measured parameters; mito^DHFR^ does not differ from cyt^DHFR^ controls. **(K)**–**(L′)** Live imaging of mito^OMM^ (magenta) alone **(K)** or co-expressed with mito-IMM^DHFR^ (green) after constitutive expression for approximately 7 days (OK371-Gal4). Control mitochondria form a continuous tubular network **(K)**. mito-IMM^DHFR^ co-expression transforms mitochondrial architecture, with the majority of organelles adopting donut-shaped structures **(L)**, **(L′)**. Donut-shaped mitochondria are visible only by live imaging and are not preserved after chemical fixation. No co-localisation of mito-IMM^DHFR^ with mito^OMM^ is detectable at this stage. Scale bars: 5 µm (overviews), 1 µm (insets). **(M)**–**(O)** Quantification of mito^OMM^-labelled mitochondria per soma in controls and mito-IMM^DHFR^ expressing animals: total area **(M)**, total surface **(N)**, and number of individual mitochondria **(O)**. **(P)**–**(R)** Quantification of individual mito^OMM^-labelled mitochondria in controls and mito-IMM^DHFR^ expressing animals: sphericity **(P)**, volume **(Q)**, and number of donut-shaped structures per soma **(R)**. **(S)**–**(V)** Acute induction of mito-IMM^DHFR^ (green) and mito^OMM^ (magenta) expression using the TARGET system (tub-Gal80^ts^; OK371-Gal4). **(S)** Control expressing mito^OMM^ alone for 15 hours shows an intact mitochondrial network. **(T)** After 15 hours of mito-IMM^DHFR^ co-expression, mito-IMM^DHFR^ co-localises with mito^OMM^ within network-organised mitochondria, but donut-shaped structures are already apparent. Scale bars: 5 µm (overviews), 1 µm (insets). **(U)** Quantification of donut-shaped structures per soma after 15 hours of expression. **(V)** Frequency of somata containing donut structures after 15 hours and 7 days of mito-IMM^DHFR^ expression. Boxplots indicate median and interquartile range; dashed grey lines represent mean control values. Data were tested for normality (Shapiro-Wilk) and analysed with parametric or non-parametric tests as appropriate, with post-hoc correction for multiple comparisons. ***p ≤ 0.001. n = 26 somata from 6 animals per genotype (E–J); n = 84 (control) and 96 (mito-IMM^DHFR^) somata from 4 animals (M–R); n = 83 (control) and 128 (mito-IMM^DHFR^) somata from 5 animals (U); n = 5 (15 h) or 4 (7 d) animals (V).

To assess mitochondrial morphology independently of import capacity, we switched to live imaging of mito^OMM^. In control animals, mito^OMM^ labelled a continuous network of interconnected tubular mitochondria within the soma (Figure 2K). Constitutive expression of mito-IMM^DHFR^ transformed mitochondrial architecture: the majority of somatic mitochondria adopted donut-shaped structures, a morphology never observed in control animals (Figure 2L, L′). Quantification revealed a nearly 50% reduction in total somatic mitochondrial area and surface, while the number of individual mitochondria was slightly increased (Figure 2M-O). At the level of individual organelles, sphericity was significantly elevated, volume was reduced, and donut-shaped structures were abundant (Figure 2P-R). Donut-shaped mitochondria were not observed after chemical fixation, either with the mito^OMM^ marker or by immunostaining for inner membrane proteins, indicating that these structures are fixation-labile and visible only by live imaging. After approximately 7 days of constitutive expression, no co-localisation of mito-IMM^DHFR^ with the altered mitochondria could be detected (Figure 2L, L′), suggesting that at this late stage, mito-IMM^DHFR^ has been degraded or cleared from the outer membrane, or that the donut structures have lost all residual import capacity. The large diameter of donut-shaped mitochondria in the soma contrasts with the small, discrete mitochondria normally present in axons (Figure 1F), raising the possibility that the morphological transformation contributes to the depletion of mitochondria from distal compartments.

To determine whether mito-IMM^DHFR^ initially co-localizes with mitochondria and to resolve the time course of morphological disruption, we acutely induced expression using the TARGET system (McGuire *et al*, 2003). After shifting larvae to the permissive temperature (29°C) for 15 hours within the third instar larval stage, we observed reliable expression of both mito-IMM^DHFR^ and mito^OMM^ (Figure 2S, T). At this early time point, mito-IMM^DHFR^ co-localised with mito^OMM^ in network-organised mitochondria, but first donut-shaped structures were already apparent. Phenotypes ranged from predominantly intact networks to numerous donut structures across individual somata (Figure 2T, Figure S1A, B). All mitochondria with donut-shaped morphology at this stage still contained both markers, confirming that mito-IMM^DHFR^ localizes to mitochondria before morphological disruption proceeds. Approximately 50% of motoneuron somata contained donut structures after 15 hours of expression, and this proportion increased to approximately 90% after 7 days (Figure 2U, V). Mitochondrial co-localisation of mito-IMM^DHFR^ was also confirmed in axons after 48 hours of expression (Figure S1C). Together, these data demonstrate that mito-IMM^DHFR^ efficiently targets mitochondria, progressively blocks protein import, and induces morphological remodelling of the mitochondrial network, with individual organelles adopting donut-shaped configurations.

### Import stress depletes functional mitochondria from synaptic terminals

Within neurons, the synaptic terminal has the highest demand for mitochondria-derived energy due to the ATP dependence of neurotransmitter release and vesicle recycling (Rangaraju et al., 2014). We therefore analysed the distribution of mitochondria at the neuromuscular junction (NMJ) of motoneurons. For all subsequent analyses of axonal and synaptic compartments, constructs were expressed in motoneurons using the OK371-Gal4 driver. The NMJ is organised into a series of synaptic boutons, each containing multiple individual release sites (active zones). In cyt^DHFR^ expressing control animals, mito^matrix^-labelled mitochondria were uniformly distributed throughout the NMJ with multiple mitochondria present within each bouton (Figure 3A, D-F). At mito^DHFR^ expressing NMJs, we observed a similar distribution and the number and size of individual mitochondria within a defined area of the NMJ remained unchanged (Figure 3B, D, E). However, the intensity of the mito^matrix^ label was significantly reduced by approximately 50%, indicating a partial impairment of mitochondrial protein import at this compartment (Figure 3B, F). In contrast, expression of mito-IMM^DHFR^ reduced the number of mitochondria at the NMJ to less than 10% of control values and all remaining mitochondria were drastically decreased in both size and staining intensity (Figure 3C-F). Analysis of NMJ area using a neuronal membrane marker revealed a significant reduction in NMJ size in mito-IMM^DHFR^ expressing animals with clear signs of neurodegeneration: in contrast to the controls where each NMJ branch terminates in a well-defined bouton, we observed distal-to-proximal fragmentation of the neuronal membrane (Figure 3C, G).

**Figure 3:**
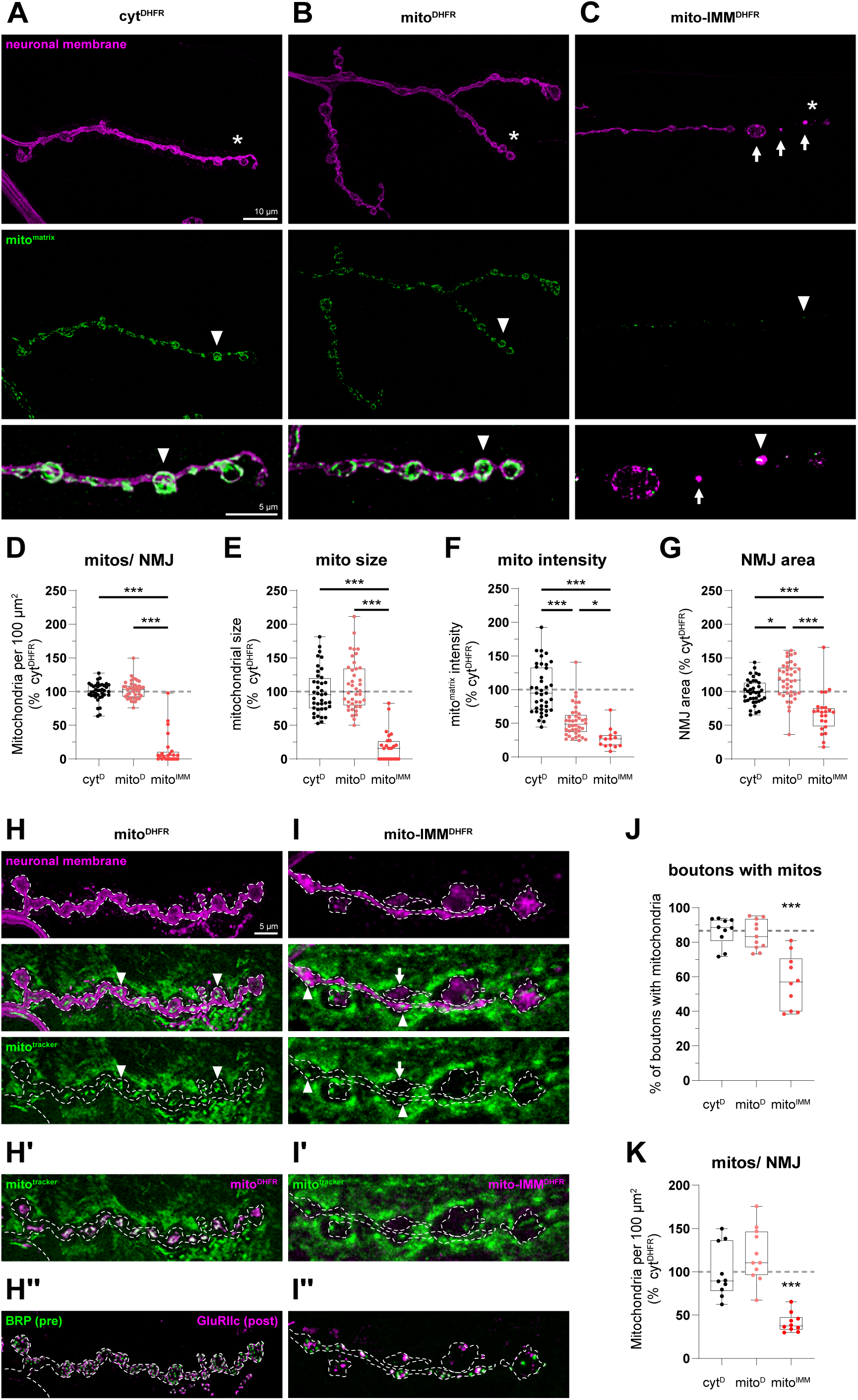
Import stress depletes mitochondria from synaptic terminals. **(A)**–**(C)** Confocal images of muscle 4 NMJs of motoneurons co-expressing mito^matrix^ with cyt^DHFR^ **(A)**, mito^DHFR^ **(B)**, or mito-IMM^DHFR^ **(C)** (OK371-Gal4). Top row: neuronal membrane (HRP, magenta). Lower rows: mito^matrix^ (HA immunostaining, green) at increasing magnification. In cyt^DHFR^ and mito^DHFR^ expressing animals, mitochondria localise to synaptic boutons throughout the NMJ (arrowheads). In mito-IMM^DHFR^ expressing animals, mitochondrial number and size are reduced, the neuronal membrane is fragmented (arrows), and mitochondria are lost from distal boutons. Asterisks indicate regions shown at higher magnification. Scale bars: 10 µm (top row), 5 µm (lower rows). **(D)**–**(G)** Quantification of mitochondrial parameters at the NMJ, normalised to cyt^DHFR^ controls. Mitochondrial density per 100 µm² **(D)**, mitochondrial size **(E)**, mito^matrix^ signal intensity **(F)**, and NMJ area **(G)**. mito^DHFR^ expression reduces mito^matrix^ intensity by approximately 50% without affecting mitochondrial number or size. All parameters are significantly reduced in mito-IMM^DHFR^ expressing animals. **(H)**–**(H″)** MitoTracker Red CMXRos labelling at muscle 6/7 NMJs of mito^DHFR^ expressing animals (OK371-Gal4). **(H)** Neuronal membrane (magenta, top) and MitoTracker (green, bottom) showing mitochondria within individual synaptic boutons (arrowheads). **(H′)** Co-imaging of MitoTracker (green) and mito^DHFR^-GFP (magenta) confirms co-localisation of the construct with MitoTracker-labelled mitochondria at the NMJ. **(H″)** The same NMJ after fixation and immunostaining for the presynaptic active zone marker Brp (green) and the postsynaptic glutamate receptor subunit GluRIIc (magenta). Dashed line indicates the NMJ boundary defined by the neuronal membrane staining in **(H)**. All boutons retain Brp-GluRIIc apposition. **(I)**–**(I″)** MitoTracker labelling at muscle 6/7 NMJs of mito-IMM^DHFR^ expressing animals. **(I)** Neuronal membrane (magenta, top) and MitoTracker (green, bottom). Mitochondria are reduced in number and size (arrowheads), and a subset of boutons is devoid of detectable mitochondria. **(I′)** MitoTracker (green) and mito-IMM^DHFR^-GFP (magenta). No GFP signal is detectable at the NMJ. **(I″)** The same NMJ after fixation and immunostaining for Brp (green) and GluRIIc (magentaA subset of boutons retains GluRIIc without Brp, indicating synaptic degeneration. Dashed line indicates the NMJ boundary. Scale bars in **(H)** apply to **(H)**–**(I″)**: 5 µm. **(J)** Percentage of boutons containing MitoTracker-positive mitochondria. cyt^DHFR^ and mito^DHFR^ expressing animals do not differ; in mito-IMM^DHFR^ expressing animals the fraction is reduced to approximately 60%. **(K)** Mitochondrial density per 100 µm² based on MitoTracker quantification. Boxplots indicate median and interquartile range; dashed grey lines represent mean control values. Data were tested for normality (Shapiro-Wilk) and analysed with parametric or non-parametric tests as appropriate, with post-hoc correction for multiple comparisons. *p ≤ 0.05, ***p ≤ 0.001. n = 39 (cyt^DHFR^), 40 (mito^DHFR^), or 23 (mito-IMM^DHFR^) NMJs from 7 animals per genotype (D–G); n = 10 (cyt^DHFR^, mito-IMM^DHFR^) or 11 (mito^DHFR^) NMJs from 6 animals per genotype (J, K).

To verify that the decreased mito^matrix^ signal at the NMJ reflects a genuine reduction in mitochondrial content rather than solely impaired import of the marker protein, we used MitoTracker Red CMXRos (Desai *et al*, 2024), a membrane potential-dependent dye that labels mitochondria independently of protein import. MitoTracker cannot be used in the soma due to high absorption by surrounding glial cells in the ventral nerve cord, but reliably labelled mitochondria at the NMJ in both the presynaptic motoneuron terminal and the postsynaptic muscle. Co-imaging of MitoTracker with the GFP-tagged constructs confirmed that the mito^DHFR^ GFP signal co-localised with MitoTracker-labelled mitochondria within individual boutons, directly confirming mitochondrial targeting at the NMJ (Figure 3H, H′). Quantification of boutons containing MitoTracker-positive mitochondria revealed no significant difference between cyt^DHFR^ and mito^DHFR^ expressing animals (Figure 3J). In contrast, only approximately 60% of boutons at mito-IMM^DHFR^ expressing NMJs still contained MitoTracker-positive mitochondria, and these were reduced in size and number, while the remaining boutons were entirely devoid of detectable mitochondria (Figure 3I, J). Quantification of mitochondrial density using this import-independent marker confirmed the significant reduction observed with mito^matrix^ (Figure 3K). In mito-IMM^DHFR^ expressing animals, no GFP signal was detectable at the NMJ (Figure 3I′), consistent with the severely reduced axonal targeting observed above (Figure 1H). Co-staining for the presynaptic active zone marker Bruchpilot (Brp) and postsynaptic glutamate receptors (GluRIIc) revealed signs of synaptic degeneration at mito-IMM^DHFR^ expressing NMJs: a subset of boutons lacked presynaptic Brp while retaining only residual postsynaptic GluRIIc clusters (Figure 3I′′), a phenotype never observed in cyt^DHFR^ or mito^DHFR^ expressing NMJs (Figure 3H′′). However, mitochondrial absence and synaptic degeneration did not perfectly coincide: boutons devoid of mitochondria frequently still retained Brp, and conversely, not all degenerated boutons had lost their mitochondria. These data confirm that mito-IMM^DHFR^ expression causes a genuine depletion of mitochondria from the presynaptic terminal and that synaptic degeneration occurs at these NMJs, but indicate that the two processes are not strictly coupled at the level of individual boutons.

### Mitochondrial protein import stress causes progressive neurodegeneration

Having established that mito-IMM^DHFR^ depletes mitochondria from the synaptic terminal (Figure 3), we next systematically characterised the impact on synaptic integrity. Synaptic degeneration at the *Drosophila* NMJ is defined by the loss of presynaptic Brp from boutons that retain postsynaptic GluRIIc, accompanied by fragmentation of the neuronal membrane (Pielage *et al*, 2005, 2008; Koch *et al*, 2008; Graf *et al*, 2011; Stephan *et al*, 2015; Mushtaq *et al*, 2022). To quantify this phenotype, we co-stained for the neuronal membrane, Brp, and GluRIIc at both muscle 4 and muscle 6/7 NMJs. In cyt^DHFR^ and mito^DHFR^ expressing control animals, NMJs at both muscle types displayed intact neuronal membranes organised into clearly identifiable boutons with precise Brp–GluRIIc apposition throughout (Figure 4A, B; Figure S2A, B). Expression of mito-IMM^DHFR^ in motoneurons caused severe synaptic degeneration at both muscle types: the neuronal membrane was fragmented, NMJ branches were thinned, and multiple boutons lacked presynaptic Brp despite the continued presence of postsynaptic GluRIIc clusters (Figure 4C; Figure S2C). This degeneration followed a distal-to-proximal pattern, indicating progressive presynaptic degeneration. Quantification revealed that more than 70% of muscle 4 NMJs and more than 80% of muscle 6/7 NMJs displayed at least one degenerated bouton (Figure 4E, G). The severity was substantial, with individual NMJs containing up to 14 degenerated boutons at muscle 4 and more than 20 at muscle 6/7 (Figure 4F, H; Figure S2E, F). Neither cyt^DHFR^ nor mito^DHFR^ expression caused detectable degeneration at either muscle type.

**Figure 4:**
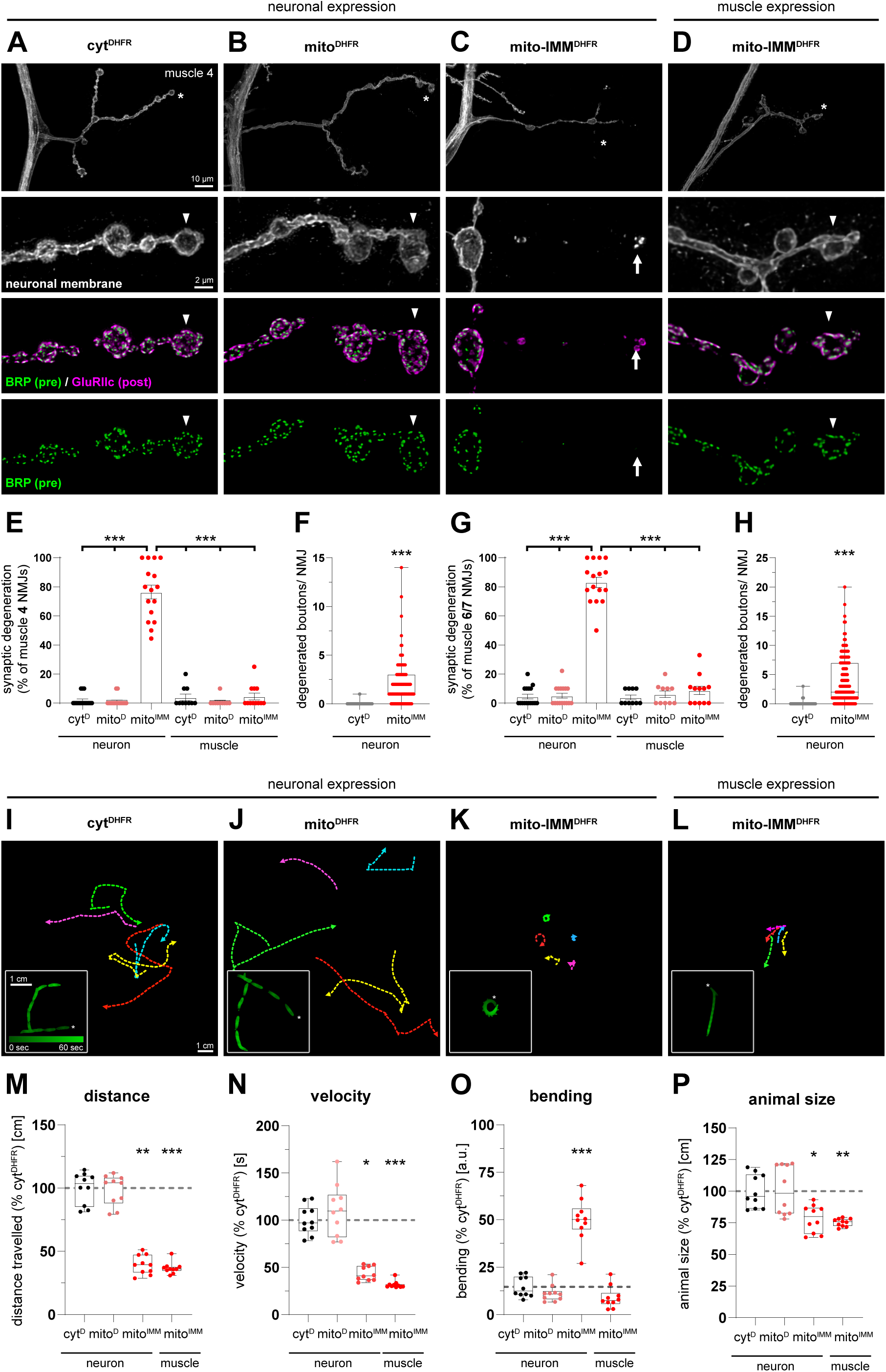
Mitochondrial protein import stress causes progressive neurodegeneration and locomotor dysfunction. **(A)**–**(D)** Confocal images of muscle 4 NMJs. Constructs were expressed in motoneurons (OK371-Gal4; **A**–**C**) or in somatic muscles (BG57-Gal4; **D**). Top row: NMJ overview with neuronal membrane (HRP, white). Lower rows at higher magnification: neuronal membrane alone, neuronal membrane (magenta), Brp (green) and GluRIIc (magenta) merge, and Brp alone. **(A)**, **(B)** cyt^DHFR^ and mito^DHFR^ expressing motoneurons display intact neuronal membranes with Brp and GluRIIc in precise apposition at all boutons (arrowheads). **(C)** Motoneuron expression of mito-IMM^DHFR^ causes fragmentation of the neuronal membrane (arrows) and loss of Brp from individual boutons despite persistent GluRIIc (arrowheads), indicating synaptic degeneration proceeding in a distal-to-proximal pattern. **(D)** Muscle expression of mito-IMM^DHFR^ does not cause synaptic degeneration; Brp and GluRIIc remain in apposition at all boutons (arrowheads). Asterisks indicate regions shown at higher magnification. Scale bars: 10 µm (overviews), 2 µm (zoom-ins). **(E)** Frequency of synaptic degeneration at muscle 4 NMJs for neuronal and muscle expression of the indicated constructs. **(F)** Severity of degeneration at muscle 4 NMJs, quantified as the number of degenerated boutons per NMJ for neuronal cyt^DHFR^ and mito-IMM^DHFR^ expression. **(G)** Frequency of synaptic degeneration at muscle 6/7 NMJs for neuronal and muscle expression. **(H)** Severity of degeneration at muscle 6/7 NMJs for neuronal cyt^DHFR^ and mito-IMM^DHFR^ expression. Only motoneuron expression of mito-IMM^DHFR^ causes significant degeneration; muscle expression does not affect synaptic integrity at either muscle type. **(I)**–**(L)** Representative locomotion traces of third instar larvae over a 2-minute observation period (top) and magnified 60-second traces colour-coded by time (bottom). **(I)** cyt^DHFR^ and **(J)** mito^DHFR^ expressing larvae crawl in straight trajectories covering the observation area. **(K)** Motoneuron expression of mito-IMM^DHFR^ causes reduced distance with pronounced lateral bending and tail-flips. **(L)** Muscle expression of mito-IMM^DHFR^ reduces crawling speed without altering directionality. Scale bars: 1 cm. **(M)**–**(P)** Quantification of locomotion parameters, normalised to cyt^DHFR^ controls. Distance travelled **(M)**, velocity **(N)**, bending **(O)**, and animal size **(P)**. Both motoneuron and muscle expression of mito-IMM^DHFR^ reduce distance, velocity, and animal size. Only motoneuron expression causes a significant increase in bending behaviour. Boxplots indicate median and interquartile range; dashed grey lines represent mean control values. Bars represent mean values with SEM. Data were tested for normality (Shapiro-Wilk) and analysed with parametric or non-parametric tests as appropriate, with post-hoc correction for multiple comparisons. *p ≤ 0.05, **p ≤ 0.01, ***p ≤ 0.001. n = 16 animals (neuronal expression) or 10 animals (muscle expression) with 10 NMJs each (E, G); n = 160 NMJs from 16 animals (neuronal expression) or 100 NMJs from 10 animals (muscle expression) (F, H); n = 10 runs with 5 animals each (M–P).

To determine whether this degeneration reflected a neuron-specific vulnerability or could also be induced by postsynaptic perturbation, we expressed mito-IMM^DHFR^ in all somatic muscles (BG57-Gal4) and analysed synaptic stability at both muscle 4 and muscle 6/7 NMJs. Muscle-driven mito-IMM^DHFR^ did not cause synaptic degeneration: the neuronal membrane remained intact, Brp and GluRIIc maintained their apposition, and neither degeneration frequency nor severity differed from control conditions at either muscle type (Figure 4D, E, G; Figure S2D–F). The only detectable difference was a minor reduction in NMJ size compared to controls (Figure 4D; Figure S2D). These data demonstrate that the progressive neurodegeneration induced by mitochondrial protein import stress is cell-autonomous to motoneurons and is not a secondary consequence of postsynaptic muscle dysfunction.

As motoneurons control the peristaltic locomotion of *Drosophila* larvae, we next tested whether import stress affected motor function. Using FIM-based locomotion tracking (Risse *et al*, 2013), we quantified crawling behaviour over two-minute observation periods. Motoneuron expression of cyt^DHFR^ and mito^DHFR^ did not impair locomotion: larvae crawled in straight trajectories covering the observation area (Figure 4I, J). Both motoneuron and muscle expression of mito-IMM^DHFR^ reduced total distance and average velocity to approximately 40% of control values (Figure 4K, L, M, N). However, the locomotion patterns differed qualitatively between the two conditions. Muscle expression reduced crawling speed but did not alter trajectory or directionality, in line with a general reduction in available muscular energy (Figure 4L, M–O). In contrast, motoneuron expression caused a pronounced increase in lateral bending and larvae displayed characteristic tail-flips (Figure 4K, O), a locomotion pattern previously associated with disrupted axonal transport and motoneuron dysfunction (Hurd & Saxton, 1996; Pielage *et al*, 2005). This behaviour, with posterior segments appearing paralyzed while anterior segments bend excessively, is consistent with a loss of coordinated innervation across body wall muscle groups. Animal size was reduced to approximately 75–80% of control values in both mito-IMM^DHFR^ conditions (Figure 4P). The qualitative difference between the two locomotion phenotypes separates a general reduction in muscular energy production, which independently confirms that the clogger impairs mitochondrial function, from the neuronal coordination deficit caused by progressive motoneuron degeneration.

### Mitochondrial protein import stress impairs synaptic transmission

To directly assess whether the structural degeneration at mito-IMM^DHFR^ expressing NMJs was accompanied by functional deficits, we performed intracellular electrophysiological recordings at muscle 6/7 NMJs. Neither cyt^DHFR^ nor mito^DHFR^ expression altered any measured parameter of basal synaptic transmission: miniature excitatory junction potential (mEJP) amplitude, evoked junction potential (EJP) amplitude, quantal content, and spontaneous release frequency were indistinguishable between the two genotypes (Figure 5A–D; Figure S3A–D). In contrast, mito-IMM^DHFR^ expression caused a significant reduction in mEJP amplitude to approximately 70% of control values. Because mEJP amplitude reflects quantal size, the amount of neurotransmitter packaged into individual synaptic vesicles, this reduction indicates impaired vesicle filling at import-stressed terminals (Figure 5A; Figure S3A). EJP amplitudes were reduced more severely, to approximately 50% of controls (Figure 5B; Figure S3B). Calculation of quantal content revealed a significant decrease, demonstrating an additional presynaptic deficit in the number of vesicles released per action potential (Figure 5C; Figure S3C). The frequency of spontaneous mEJPs was reduced by approximately 40% (Figure 5D; Figure S3D), in line with the loss of presynaptic active zones observed at these NMJs (Figure 4).

**Figure 5:**
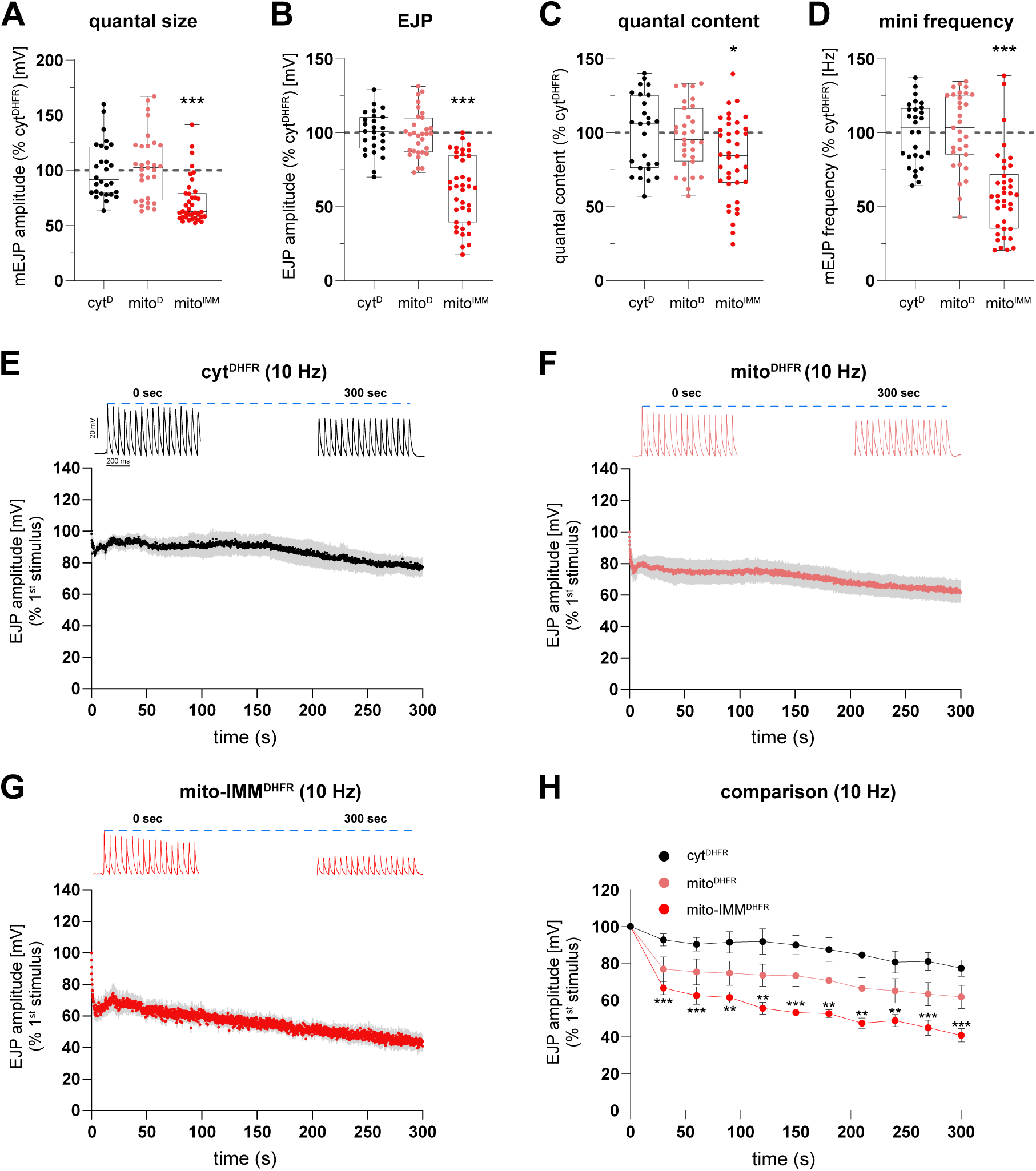
Mitochondrial protein import stress impairs synaptic transmission. **(A)**–**(D)** Intracellular electrophysiological recordings at muscle 6/7 NMJs of motoneurons expressing cyt^DHFR^, mito^DHFR^, or mito-IMM^DHFR^ (OK371-Gal4). All values are normalised to cyt^DHFR^ controls. **(A)** Miniature excitatory junction potential (mEJP) amplitude. **(B)** Evoked excitatory junction potential (EJP) amplitude. **(C)** Quantal content. **(D)** Spontaneous mEJP frequency. In mito-IMM^DHFR^ expressing animals, mEJP amplitude, EJP amplitude, quantal content, and mEJP frequency are all significantly reduced. mito^DHFR^ does not differ from cyt^DHFR^ controls. **(E)**–**(G)** EJP amplitude during continuous 10 Hz stimulation for 300 seconds, normalised to the first stimulus. Representative traces at 0 s and 300 s are shown above each graph. Mean (line) and range (shading) are plotted over time; dashed blue line indicates 100% of initial amplitude. **(E)** cyt^DHFR^ controls show mild decline, stabilising above 80%. **(F)** mito^DHFR^ expressing animals show a modest initial drop, stabilising around 70–80%. **(G)** mito-IMM^DHFR^ expressing animals show progressive decline throughout the stimulation period, reaching approximately 40% by 300 seconds. **(H)** Comparison of EJP amplitude over time during 10 Hz stimulation for all three genotypes. mito-IMM^DHFR^ expressing animals are significantly reduced from the onset and throughout the stimulation period. Boxplots indicate median and interquartile range; dashed grey lines represent mean control values. Data were tested for normality (Shapiro-Wilk) and analysed with parametric or non-parametric tests as appropriate, with post-hoc correction for multiple comparisons. *p ≤ 0.05, **p ≤ 0.01, ***p ≤ 0.001. n = 28 (cyt^DHFR^), 31 (mito^DHFR^), or 38 (mito-IMM^DHFR^) recordings from 10 (cyt^DHFR^, mito^DHFR^) or 12 (mito-IMM^DHFR^) animals (A–D); n = 6 (cyt^DHFR^), 10 (mito^DHFR^), or 9 (mito-IMM^DHFR^) animals (E–H).

To test the capacity for sustained neurotransmitter release, we challenged NMJs with continuous 10 Hz stimulation for 300 seconds. In cyt^DHFR^ expressing controls, EJP amplitudes showed only a mild decline and stabilised above 80% of the initial response (Figure 5E). mito^DHFR^ expressing animals displayed a modest initial drop to approximately 80%, stabilizing around 70% of the initial amplitude; this trend did not reach significance (Figure 5F, H). In mito-IMM^DHFR^ expressing animals, EJP amplitudes were already reduced at the onset of stimulation and declined progressively throughout the protocol, reaching approximately 40% of the initial response by 300 seconds (Figure 5G, H). These data demonstrate that mitochondrial protein import stress impairs both initial and sustained synaptic transmission at the NMJ.

### Mitochondrial absence at the synapse does not cause neurodegeneration

The degeneration observed in mito-IMM^DHFR^ expressing motoneurons could, in principle, result from the depletion of mitochondria from the presynaptic terminal rather than from the import block itself. To distinguish between these possibilities, we analysed *miro* mutant animals (*miro*^sd32^/^b682^), in which mitochondrial transport to the synapse is abolished (Guo *et al*, 2005). Miro is a mitochondrial Rho-GTPase required for kinesin-dependent anterograde transport of mitochondria; in its absence, mitochondria accumulate in neuronal somata and are absent from axons and synaptic terminals. Visualization of mitochondria using mito^matrix^ confirmed that in *miro* mutant motoneurons, mitochondria formed dense somatic accumulations without discernible network structure and were absent from axons (Figure 6A, B). Mitochondria were completely eliminated from both muscle 6/7 (Figure 6C, D, E) and muscle 4 NMJs. Despite this complete depletion of mitochondria, which exceeds the mitochondrial reduction observed in mito-IMM^DHFR^ expressing animals, we only observed minor alterations to NMJ morphology with no significant reduction in NMJ size (Figure 6F). The principal morphological change was the emergence of small satellite boutons from main NMJ branches (Figure 6D), a phenotype previously reported for *miro* mutants and attributed to altered presynaptic microtubule organization (Guo *et al*, 2005; Russo *et al*, 2009).

**Figure 6:**
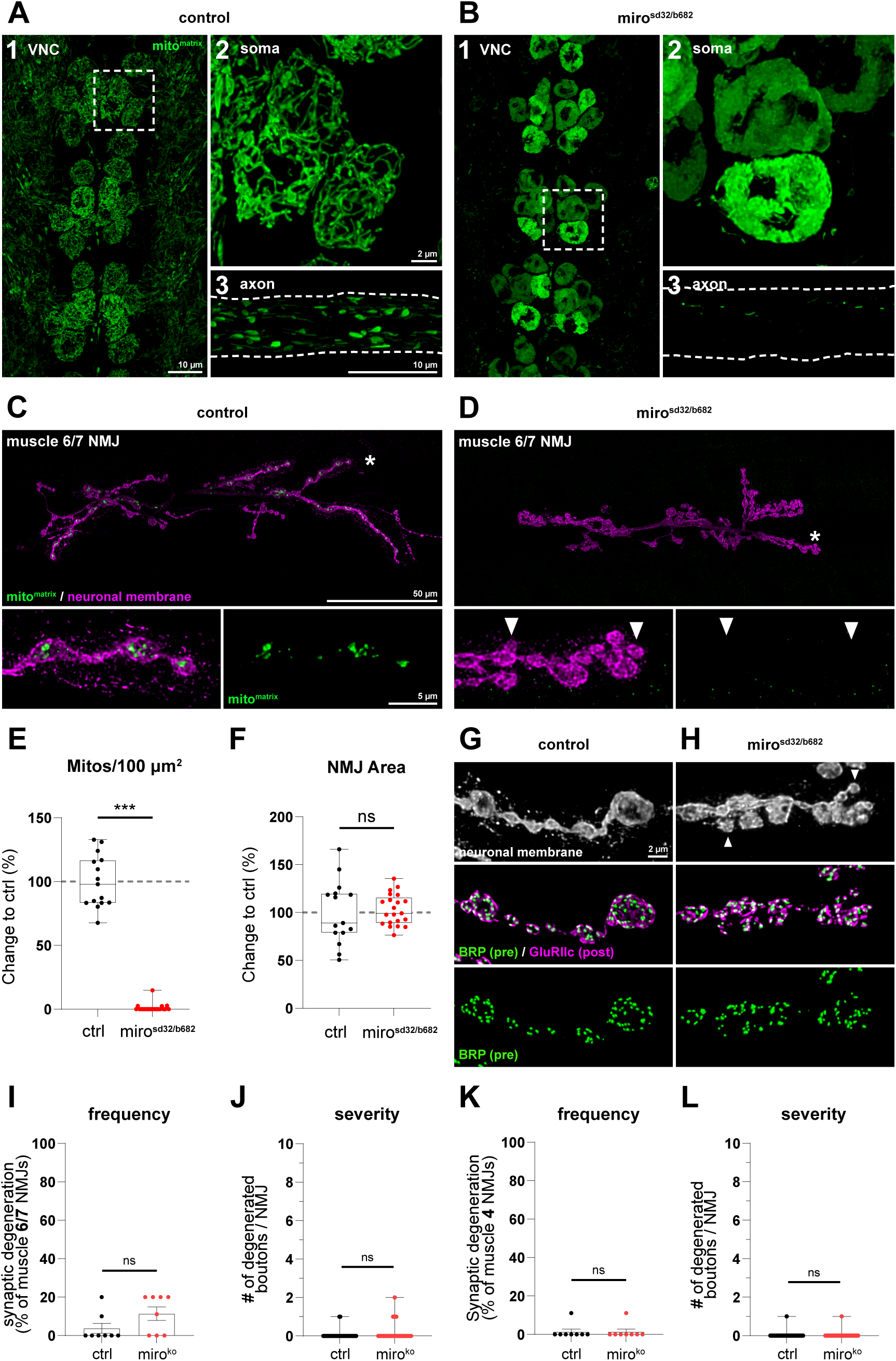
Mitochondrial absence at the synapse does not cause neurodegeneration. **(A)**, **(B)** Mitochondrial distribution in control **(A)** and *miro*^sd32/b682^ mutant **(B)** motoneurons (OK371-Gal4). mito^matrix^ (green) labels the mitochondrial network. Sub-panels show: (1) ventral nerve cord overview, (2) individual soma at higher magnification, (3) motoneuron axons. In controls, mitochondria form an interconnected network in the soma and distribute evenly through axons. In *miro* mutants, mitochondria accumulate in dense somatic clusters without discernible network structure and are nearly completely absent from axons. Dashed boxes indicate regions shown at higher magnification. Scale bars: 10 µm (VNC), 2 µm (soma), 10 µm (axons). **(C)**, **(D)** Confocal images of muscle 6/7 NMJs in control **(C)** and *miro*^sd32/b682^ mutant **(D)** animals. mito^matrix^ (green) and neuronal membrane (HRP, magenta). In controls, mitochondria localise to synaptic boutons throughout the NMJ. In *miro* mutants, mitochondria are absent from the NMJ; small satellite boutons emerge from main branches (arrowheads). Asterisks indicate regions shown at higher magnification. Scale bars: 50 µm (overviews), 5 µm (zoom-ins). **(E)** Mitochondrial density per 100 µm² at muscle 6/7 NMJs, normalised to controls. No mitochondria are detected at *miro* mutant NMJs. **(F)** NMJ area, normalised to controls. NMJ area is not significantly changed in *miro* mutants despite the complete absence of presynaptic mitochondria. **(G)**, **(H)** Confocal images of muscle 6/7 NMJs in control **(G)** and *miro*^sd32/b682^ mutant **(H)** animals stained for the neuronal membrane (HRP, white), Brp (green), and GluRIIc (magenta), with Brp shown separately. In controls, Brp and GluRIIc are in precise apposition at all boutons. In *miro* mutants, Brp-GluRIIc apposition is maintained throughout the NMJ despite the emergence of satellite boutons (arrowhead). **(I)** Frequency of synaptic degeneration at muscle 6/7 NMJs. **(J)** Severity of degeneration at muscle 6/7 NMJs. **(K)** Frequency of synaptic degeneration at muscle 4 NMJs. **(L)** Severity of degeneration at muscle 4 NMJs. No significant increase in degeneration frequency or severity is detected at either muscle type. Boxplots indicate median and interquartile range; bars represent mean values with SEM; dashed grey lines represent mean control values. Data were tested for normality (Shapiro-Wilk) and analysed with parametric or non-parametric tests as appropriate, with post-hoc correction for multiple comparisons. ns, p ≥ 0.05, ***p ≤ 0.001. n = 15 (control) or 21 (*miro*^sd32/b682^) NMJs from 6 animals each (E, F); n = 8 animals with 10 NMJs each (I–L).

To directly assess synaptic stability, we co-stained for the neuronal membrane, Brp, and GluRIIc. In *miro* mutant animals, the neuronal membrane remained intact and all presynaptic Brp puncta were precisely apposed by postsynaptic GluRIIc clusters at both muscle 6/7 (Figure 6G, H) and muscle 4 NMJs. Quantification confirmed no significant increase in degeneration frequency or severity at either muscle type (Figure 6I–L), consistent with the absence of synaptic degeneration previously noted for *miro* mutant NMJs (Massaro *et al*, 2009). These data demonstrate that the complete absence of mitochondria from the presynaptic terminal is not sufficient to induce neurodegeneration at the *Drosophila* NMJ. The progressive degeneration observed in mito-IMM^DHFR^ expressing motoneurons therefore cannot be explained by mitochondrial depletion alone but must instead arise from the import block itself or its cellular consequences.

### Mitochondrial protein import stress induces a conserved and neuron-specific transcriptomic response

Having demonstrated that sustained motoneuron expression of mito-IMM^DHFR^ causes synaptic degeneration (Figure 4), we next asked how this phenotype develops over time and whether it is accompanied by a transcriptional stress response. We used the TARGET system (tub-Gal80^ts^; n-Syb-Gal4) to express the clogger (UAS-mito-IMM^DHFR^) for either one, two or three days prior to analysis, collecting only wandering third instar larvae to ensure that developmental stage was matched across all conditions. Degeneration, defined as loss of the presynaptic active zone marker Brp from boutons that retain postsynaptic GluRIIC, was absent at Day 1, first detectable at Day 2 and became significant by Day 3 of mito-IMM^DHFR^ expression (Figure 7A, B). We then performed RNA-seq on dissected larval brains along this same timeline, comparing animals after one, two or three days of clogger expression to uninduced controls (Day 0). This analysis revealed a progressive transcriptomic response that increased in both the number of affected genes and the magnitude of expression changes with each additional day of import stress.

**Figure 7:**
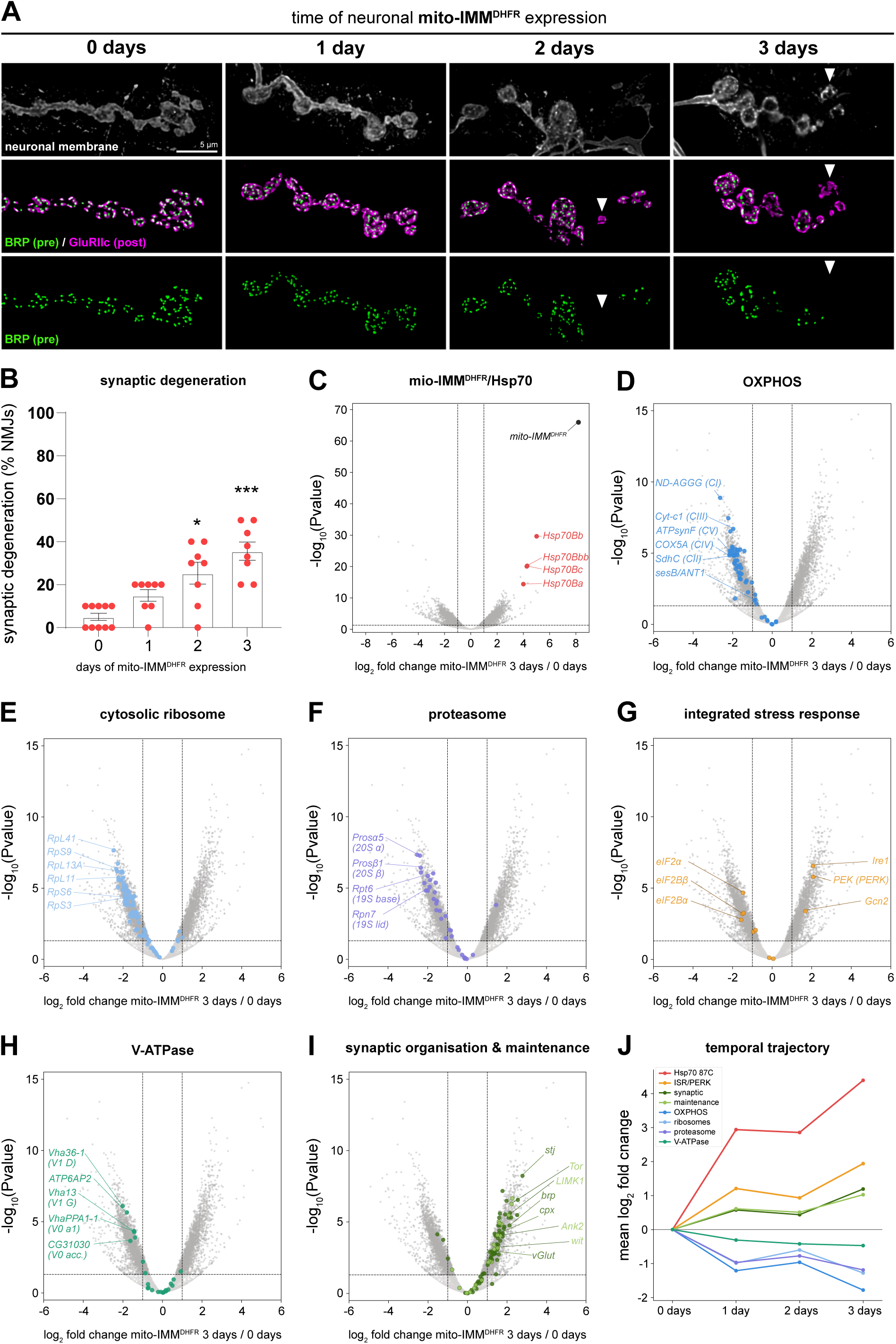
Import stress induces a conserved and neuron-specific transcriptomic response. **(A)** Confocal images of muscle 6/7 NMJs after 0, 1, 2, or 3 days of mito-IMM^DHFR^ expression prior to analysis at the wandering third instar larval stage, using the TARGET system (n-syb-Gal4; tub-Gal80^ts^). All animals are of the same developmental stage but differ in the duration of mito-IMM^DHFR^ expression. Neuronal membrane (HRP, white), Brp (green) and GluRIIc (magenta) merge, and Brp alone are shown for each condition. Degeneration, defined as loss of Brp from boutons that retain GluRIIc, is absent after 0 and 1 day of expression, first apparent after 2 days (arrowheads), and pronounced after 3 days. Scale bar: 5 µm. **(B)** Quantification of synaptic degeneration frequency. Degeneration is significantly increased after 2 days and further elevated after 3 days of mito-IMM^DHFR^ expression. **(C)** Volcano plot of differential gene expression after 3 days of mito-IMM^DHFR^ expression compared to uninduced controls (0 days). The transgene mito-IMM^DHFR^ is the most strongly upregulated gene. The four *Hsp70* 87C locus genes (*Hsp70Ba*, *Hsp70Bb*, *Hsp70Bbb*, *Hsp70Bc*) are among the most significantly induced endogenous transcripts. Dashed lines indicate significance thresholds. **(D)**–**(I)** Pathway-specific volcano plots highlighting gene sets of interest within the 3 days dataset. Coloured points indicate genes belonging to the indicated category; grey points represent all other detected genes. Selected genes are labelled. **(D)** Nuclear-encoded oxidative phosphorylation (OXPHOS) subunits of Complexes I–V and the adenine nucleotide translocase *sesB*; the majority are significantly downregulated. **(E)** Cytosolic ribosomal protein genes (40S and 60S subunits); broadly downregulated. **(F)** Proteasome subunits (20S catalytic core and 19S regulatory particle); broadly downregulated. **(G)** Integrated stress response components: the eIF2α kinases *PEK*/PERK and *Gcn2* are upregulated; the ER stress sensor *Ire1* is induced; eIF2α and eIF2B subunits are repressed. **(H)** Vacuolar H^+^-ATPase (V-ATPase) subunits of the V1 and V0 subcomplexes, including the accessory subunit ATP6AP2 and the neuron-specific regulator CG31030; broadly downregulated. **(I)** Synaptic organisation and maintenance genes, including active zone scaffold components, synaptic vesicle machinery, cytoskeletal regulators, and kinases previously shown to oppose synaptic degeneration; broadly upregulated. **(J)** Temporal trajectory of mean log_2_ fold change for the indicated gene categories across 0 to 3 days of mito-IMM^DHFR^ expression. Metabolic and biosynthetic pathways (OXPHOS, ribosomes, proteasome, V-ATPase) are progressively repressed, while *Hsp70* 87C, ISR/PERK components, and synaptic maintenance genes are progressively upregulated. Bars represent mean values with SEM (B). Data were tested for normality (Shapiro-Wilk) and analysed with parametric or non-parametric tests as appropriate, with post-hoc correction for multiple comparisons (B). *p ≤ 0.05, ***p ≤ 0.001. n = 10 (0 days) or 8 (1–3 days) animals with 10 NMJs each (A, B). RNA-seq was performed on dissected larval brains; n = 2 (0 days) or 3 (1–3 days) biological replicates per condition (C–J).

The most prominently induced transcripts encoded *Hsp70*, the major stress-inducible chaperone conserved across eukaryotes. In yeast, *Hsp70* induction via the heat shock transcription factor Hsf1 is the defining feature of the mitoprotein-induced stress response (Boos *et al*, 2019). The *Drosophila* genome contains six *Hsp70* gene copies distributed across two chromosomal clusters: two genes at the 87A locus (Hsp70Aa, Hsp70Ab) and four at the 87C locus (Hsp70Ba, Hsp70Bb, Hsp70Bbb, Hsp70Bc), with the related gene *Hsp68* at a separate position (Gong & Golic, 2006). Although all six genes encode nearly identical proteins and are induced by heat shock, the two loci differ in their regulatory elements and show tissue– and context-dependent transcriptional differences (Lakhotia & Prasanth, 2002). Because the temperature shift to 29°C required for TARGET-mediated transgene induction constitutes a mild heat stress, we exploited this dual-locus organisation to distinguish thermal from import stress-related *Hsp70* induction. The 87A locus genes and *Hsp68* were strongly induced at Day 1 but returned to baseline expression by Day 3, consistent with transient heat acclimation (Figure S4a). In contrast, all four 87C locus genes showed sustained and progressive induction from Day 1 through Day 3, reaching four– to five-log_2_ fold upregulation (Figure 7C, Figure S4a). The heat shock transcription factor *Hsf* was itself mildly upregulated (log_2_FC = 0.9, p < 0.01), supporting ongoing transcriptional drive. This temporal separation demonstrates that the Day 3 transcriptomic profile is dominated by the import stress response rather than by residual heat shock effects, and identifies the 87C cluster as the *Drosophila* counterpart of the conserved mitoprotein-induced *Hsp70* response described in yeast (Boos *et al*, 2019). We therefore focused subsequent pathway analyses on the Day 3 dataset. The selectivity of the chaperone response extended beyond *Hsp70*: while the 87C cluster was massively induced, small heat shock proteins, Hsp90 and the CCT/TRiC chaperonin complex were all repressed (Figure S4b), indicating specific remodelling of the chaperone network rather than a general stress response.

Import stress was accompanied by broad repression of genes encoding core cellular machinery. Nuclear-encoded subunits of all five oxidative phosphorylation complexes were reduced, including representatives of Complex I (NADH dehydrogenase), Complex II (succinate dehydrogenase), Complex III (cytochrome bc1), Complex IV (cytochrome c oxidase) and Complex V (ATP synthase), as well as the adenine nucleotide translocase *sesB* (Figure 7D). Cytosolic ribosomal protein genes were similarly affected, with 66 of 86 detected subunits of the 40S or 60S ribosome significantly depleted (Figure 7E), as were mitochondrial ribosomal protein genes (Figure S4c). Proteasome subunits of both the 20S catalytic core and the 19S regulatory particle were also repressed (Figure 7F). The repression of OXPHOS and ribosomal genes is consistent with the conserved transcriptional response to mitochondrial protein import stress in yeast, where these pathways are coordinately downregulated (Boos *et al*, 2019). The proteasome response, however, diverges: in yeast, the transcription factor *Rpn4* drives proteasome upregulation as part of the mitoprotein-induced stress response, but *Drosophila* lacks an *Rpn4* orthologue and instead shows proteasome repression, likely as part of a broader translational attenuation programme. All three pathways showed progressive decline from Day 1 through Day 3, with the magnitude of repression increasing at each timepoint (Figure 7J). Genes encoding antioxidant defence proteins were also broadly repressed, including the superoxide dismutases *Sod1* and *Sod2*, multiple glutathione S-transferases, and the peroxiredoxin Prx5 (Figure S4d).

In addition to the metabolic repression, the transcriptomic data pointed toward activation of the integrated stress response (ISR), a conserved signalling pathway that attenuates global translation through phosphorylation of the translation initiation factor eIF2α (Hetz & Mollereau, 2014). *Drosophila* encodes two eIF2α kinases: *PEK*, the orthologue of the ER-resident kinase PERK, and the cytosolic kinase *Gcn2*. Both were transcriptionally upregulated during import stress (*PEK* log_2_FC = 2.07, *Gcn2* log_2_FC = 1.68; Figure 7G). The ER stress sensor *Ire1*, which operates through a distinct mechanism involving Xbp1 mRNA splicing, was similarly induced (log_2_FC = 2.07), indicating ER stress pathway activation alongside the ISR. In parallel, the eIF2α transcript and two subunits of its recycling factor eIF2B were significantly reduced (Figure 7G), a pattern that would reinforce translational attenuation. The convergent upregulation of both eIF2α kinases with repression of eIF2B components identified the ISR as a candidate neuroprotective pathway to be tested functionally.

Beyond these conserved stress responses, import-stressed motoneurons showed transcriptional changes with direct relevance to the synaptic phenotypes. Multiple subunits of the vacuolar H+-ATPase (V-ATPase) were significantly reduced, including V1 and V0 subcomplexes as well as the accessory subunit ATP6AP2 and the neuron-specific regulator CG31030/VhaAC45RP (Figure 7H). The V-ATPase is the proton pump responsible for acidifying synaptic vesicles, thereby generating the electrochemical gradient that drives vesicular neurotransmitter loading (Forgac, 2007). At the *Drosophila* NMJ, loss of CG31030 impairs synaptic vesicle acidification and reduces quantal size (Dulac *et al*, 2021), and the V0 subunit Vha100-1 is required for synaptic vesicle exocytosis (Hiesinger *et al*, 2005). Reduced V-ATPase expression would therefore be expected to impair vesicle filling and diminish quantal size, consistent with the electrophysiological deficits we observed at import-stressed NMJs (Figure 5).

Despite this broad metabolic and biosynthetic shutdown, the transcriptomic response also included widespread upregulation of synaptic genes (Figure 7I). Using gene ontology-based categories for synaptic function (active zone scaffold components, synaptic vesicle fusion and recycling machinery, endocytic factors, ion channels, and synaptic adhesion molecules), we found that 48 of 66 detected genes were significantly upregulated, with only three significantly reduced. In parallel, we assembled a manually curated set of 22 genes with established roles in *Drosophila* NMJ maintenance, drawing on published genetic studies of synapse stability (Pielage *et al*, 2005, 2008; Koch *et al*, 2008; Eaton & Davis, 2005; Enneking *et al*, 2013; Mushtaq *et al*, 2022; Stephan *et al*, 2015; Graf *et al*, 2011). Of these, 15 were significantly upregulated, including spectrin-ankyrin cytoskeletal components, BMP signalling receptors, and kinases previously shown to oppose synaptic degeneration (Figure 7I). The upregulation of both synaptic function and synapse maintenance genes indicates that import-stressed motoneurons direct a transcriptional response toward preserving synaptic integrity, even as core metabolic pathways are repressed.

The temporal trajectory of these transcriptional changes reveals two opposing programmes operating in parallel (Figure 7J, Figure S4d). Core metabolic and biosynthetic pathways, including OXPHOS, ribosomal genes, proteasome and V-ATPase, are progressively repressed from Day 1 through Day 3. Synaptic function and maintenance genes, in contrast, are broadly upregulated over the same period. The import stress response in *Drosophila* motoneurons thus shares the conserved metabolic shutdown characterised in yeast (Boos *et al*, 2019), but additionally activates neuron-specific programmes directed at maintaining synaptic integrity. Among the stress-responsive pathways, the coordinate upregulation of the ISR kinase *PEK*/PERK and the transcriptional changes in the eIF2 pathway identified PERK signalling as a candidate whose neuroprotective potential could be tested directly. We therefore directly tested whether PERK activity opposes synaptic degeneration during mitochondrial protein import stress.

### PERK-eIF2α signalling opposes neurodegeneration during mitochondrial protein import stress

The transcriptomic analysis identified the ISR kinase *PEK*/PERK as upregulated during mitochondrial protein import stress. PERK phosphorylates eIF2α to attenuate global translation, and the functional consequences of this response for neuronal survival are context-dependent, with PERK activity being neuroprotective in some disease models and neurodegenerative in others (Wang *et al*, 2011; Tsuyama *et al*, 2017). To determine the consequences of altered PERK levels in motoneurons, we first expressed a PERK-directed RNAi construct using OK371-Gal4. Motoneuron-specific PERK knockdown did not affect NMJ morphology: the neuronal membrane remained intact, Brp and GluRIIC maintained their apposition, and neither degeneration frequency nor severity differed from control animals at muscle 6/7 or muscle 4 NMJs (Figure S5A, B, D–G). Loss of PERK signalling in the absence of import stress is therefore not required for synaptic maintenance under baseline conditions.

In contrast, motoneuron-specific overexpression of PERK (Malzer *et al*, 2010) caused pronounced synaptic degeneration. NMJs displayed thinned branches with individual boutons lacking Brp despite persistent GluRIIC (Figure S5C, arrowheads). Approximately 45% of muscle 6/7 NMJs contained at least one degenerated bouton, with an average severity of 2–3 affected boutons per NMJ (Figure S5D, E). Degeneration was also significant at muscle 4 NMJs, albeit at lower frequency and severity (Figure S5F, G). PERK overexpression, in the absence of mitochondrial protein import stress, is thus sufficient to cause synaptic degeneration in motoneurons. The endogenous upregulation of PERK observed during import stress could therefore represent a protective response that mitigates damage, or alternatively, the resulting translational repression could itself contribute to the degenerative process.

To distinguish between these possibilities, we fed larvae the selective PERK inhibitor GSK2606414 (GSK) (Moreno *et al*, 2013). Treatment of cyt^DHFR^ expressing control animals with GSK did not affect NMJ morphology, degeneration frequency or severity at either muscle 4 or muscle 6/7 (Figure 8A, A′, D, E; Figure S6), consistent with the absence of a phenotype upon PERK knockdown (Figure S5). In contrast, GSK treatment of mito-IMM^DHFR^ expressing animals significantly increased both the frequency and severity of synaptic degeneration compared to mock-treated mito-IMM^DHFR^ controls (Figure 8B, B′, D, E; Figure S6). The degeneration was not only more frequent but also qualitatively more severe: GSK-treated mito-IMM^DHFR^ animals showed a substantial proportion of completely eliminated NMJs, muscles that had lost all presynaptic innervation, a phenotype that was rarely observed with import stress alone (Figure 8F). Pharmacological inhibition of PERK thus converts a partially degenerative phenotype into severe synaptic elimination, including complete loss of presynaptic innervation at affected NMJs, indicating that endogenous PERK signalling normally limits the rate and extent of neurodegeneration during mitochondrial protein import stress. In the absence of this protective response, import-stressed motoneurons rapidly progress to complete loss of synaptic connectivity.

**Figure 8:**
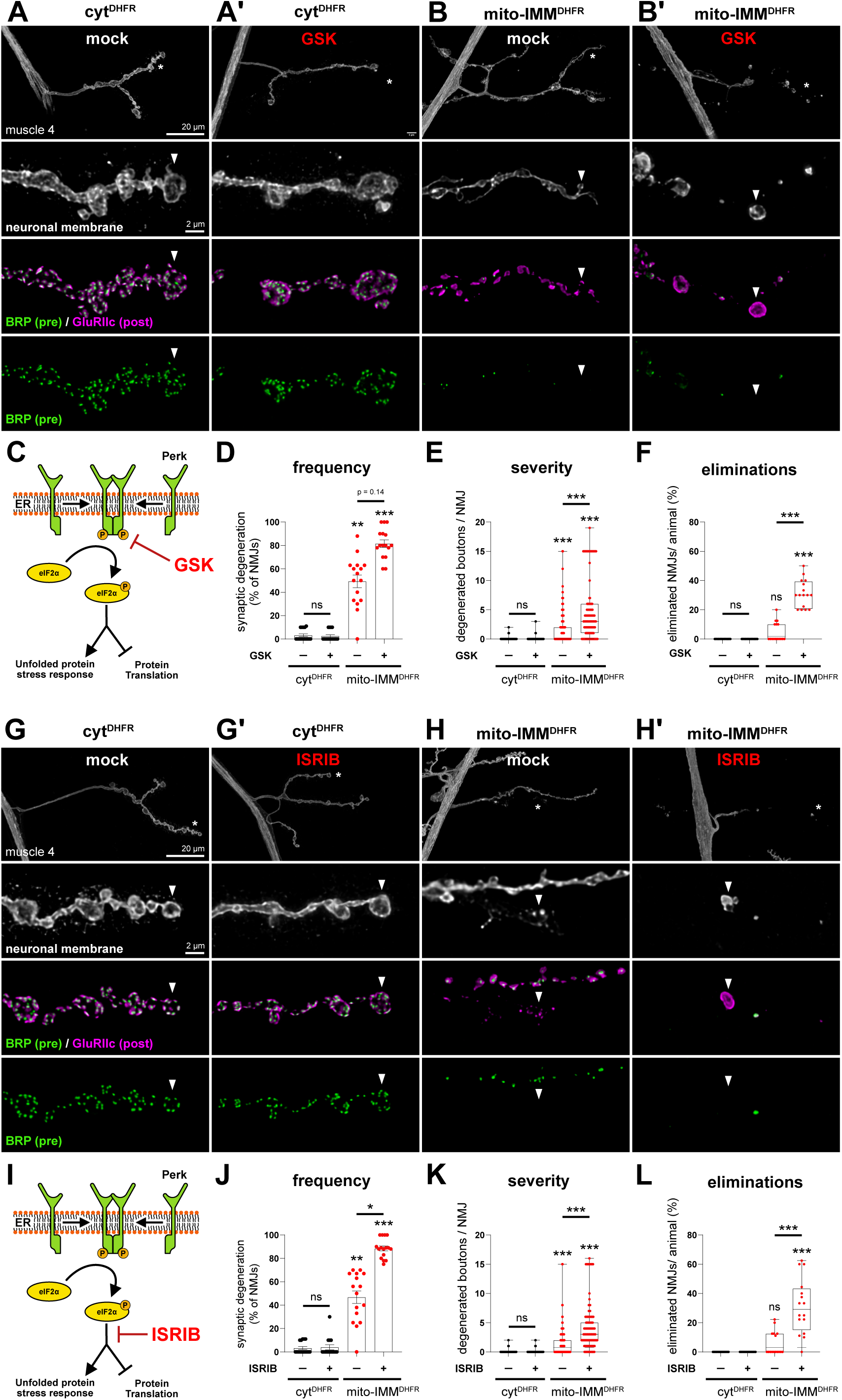
PERK-eIF2α signalling opposes neurodegeneration during mitochondrial protein import stress. **(A)**–**(B′)** Confocal images of muscle 4 NMJs of motoneurons expressing cyt^DHFR^ (**A**, **A′**) or mito-IMM^DHFR^ (**B**, **B′**) (OK371-Gal4), reared on food containing DMSO (mock, **A**, **B**) or the PERK inhibitor GSK2606414 (GSK, **A′**, **B′**). Top row: neuronal membrane (HRP, white) overview. Lower rows at higher magnification: neuronal membrane, Brp (green) and GluRIIc (magenta) merge, and Brp alone. cyt^DHFR^ expressing animals show intact NMJs in both mock **(A)** and GSK **(A′)** conditions, with Brp and GluRIIc in precise apposition at all boutons (arrowheads). mito-IMM^DHFR^ expression causes synaptic degeneration in mock-treated animals **(B)**, with loss of Brp from individual boutons despite persistent GluRIIc (arrowheads). GSK treatment of mito-IMM^DHFR^ expressing animals **(B′)** enhances degeneration and causes complete NMJ elimination, in which synaptic bouton remnants are no longer connected by the neuronal membrane. Asterisks indicate regions shown at higher magnification. Scale bars: 20 µm (overview), 2 µm (zoom-ins). **(C)** Schematic of the PERK signalling pathway. Upon activation by unfolded proteins, PERK dimerises and phosphorylates eIF2α, attenuating global translation. GSK inhibits PERK kinase activity. **(D)** Frequency of synaptic degeneration. GSK treatment has no effect on cyt^DHFR^ expressing animals but significantly increases degeneration frequency in mito-IMM^DHFR^ expressing animals. **(E)** Severity of degeneration, quantified as the number of degenerated boutons per NMJ. GSK treatment significantly increases severity in mito-IMM^DHFR^ expressing animals. **(F)** Percentage of eliminated NMJs, defined as near complete loss of Brp and complete fragmentation of the neuronal membrane. GSK treatment of mito-IMM^DHFR^ expressing animals causes a substantial proportion of NMJ eliminations. Boxplots indicate median and interquartile range; bars represent mean values with SEM. Data were tested for normality (Shapiro-Wilk) and analysed with parametric or non-parametric tests as appropriate, with post-hoc correction for multiple comparisons. ns, p ≥ 0.05, **p ≤ 0.01, ***p ≤ 0.001. n = 16 animals with 10 NMJs each per condition (D, F); n = 160 NMJs from 16 animals per condition (E). **(G)**–**(H′)** Confocal images of muscle 4 NMJs of motoneurons expressing cyt^DHFR^ (**G**, **G′**) or mito-IMM^DHFR^ (**H**, **H′**) (OK371-Gal4), reared on food containing DMSO (mock, **G**, **H**) or ISRIB (**G′**, **H′**). Top row: neuronal membrane (HRP, white) overview. Lower rows at higher magnification: neuronal membrane, Brp (green) and GluRIIc (magenta) merge, and Brp alone. cyt^DHFR^ expressing animals show intact NMJs in both mock **(G)** and ISRIB **(G′)** conditions, with Brp and GluRIIc in precise apposition at all boutons (arrowheads). mito-IMM^DHFR^ expression causes synaptic degeneration in mock-treated animals **(H)**. ISRIB treatment of mito-IMM^DHFR^ expressing animals **(H′)** enhances degeneration and causes complete NMJ elimination, in which synaptic bouton remnants are no longer connected by the neuronal membrane. Asterisks indicate regions shown at higher magnification. Scale bars: 20 µm (overview), 2 µm (zoom-ins). **(I)** Schematic of the eIF2α signalling pathway. ISRIB stabilises eIF2B in its active decameric form, restoring translation initiation downstream of eIF2α phosphorylation and thereby overriding PERK-mediated translational attenuation. **(J)** Frequency of synaptic degeneration. ISRIB has no effect on cyt^DHFR^ expressing animals but significantly increases degeneration frequency in mito-IMM^DHFR^ expressing animals. **(K)** Severity of degeneration. ISRIB treatment significantly increases severity in mito-IMM^DHFR^ expressing animals. **(L)** Percentage of eliminated NMJs, defined as near complete loss of Brp and complete fragmentation of the neuronal membrane. ISRIB treatment of mito-IMM^DHFR^ expressing animals causes a substantial proportion of NMJ eliminations. Boxplots indicate median and interquartile range; bars represent mean values with SEM. Data were tested for normality (Shapiro-Wilk) and analysed with parametric or non-parametric tests as appropriate, with post-hoc correction for multiple comparisons. ns, p ≥ 0.05, *p ≤ 0.05, **p ≤ 0.01, ***p ≤ 0.001. n = 16 animals with 10 NMJs each per condition (J, L); n = 160 NMJs from 16 animals per condition (K).

GSK inhibits PERK directly, but PERK is only one of several kinases that phosphorylate eIF2α. To test whether the protective effect operates through eIF2α-mediated translational control rather than a PERK-specific function, we used ISRIB, a small molecule that stabilises eIF2B in its active decameric form and thereby restores translation initiation downstream of all eIF2α kinases (Sidrauski *et al*, 2015). ISRIB treatment of cyt^DHFR^ expressing control animals did not cause synaptic degeneration at either muscle 4 or muscle 6/7 (Figure 8G, G′, J, K; Figure S6). In mito-IMM^DHFR^ expressing animals, ISRIB significantly increased degeneration frequency and severity and caused NMJ eliminations (Figure 8H, H′, J–L; Figure S6), phenocopying the effect of GSK. The convergence of two pharmacological approaches that target distinct nodes of the same pathway, PERK kinase activity upstream and eIF2B-mediated translation reinitiation downstream, demonstrates that the protective mechanism operates through eIF2α phosphorylation and the resulting attenuation of translation. PERK-dependent translational control via the integrated stress response thus serves as an endogenous neuroprotective programme that opposes synaptic degeneration during mitochondrial protein import stress.

## Discussion

Here, we demonstrate that mitochondrial protein import stress causes progressive synaptic degeneration in *Drosophila* motoneurons, and that PERK-dependent translational control through the integrated stress response opposes this process. Using the yeast clogger system adapted to this context with temporal control, we show that sustained blockade of the TOM-TIM23 import channel triggers a degenerative cascade from somatic mitochondrial remodelling through synaptic mitochondrial depletion to progressive structural neurodegeneration. This degeneration is mechanistically distinct from mitochondrial absence, because *miro* mutant neurons that completely lack presynaptic mitochondria do not degenerate (Guo *et al*, 2005; Massaro *et al*, 2009), and instead arises from import stress itself: compromised mitochondrial function at the synapse coupled with the predicted cytosolic accumulation of non-imported precursors. The transcriptomic response shares core features with the yeast programme but additionally activates the integrated stress response, and pharmacological experiments demonstrate that this pathway is functionally neuroprotective. PERK-dependent translational attenuation during import stress may serve a function analogous to proteasomal clearance in yeast, reducing the cytosolic precursor burden by limiting new precursor production. This positions PERK as protective when the upstream insult is mitochondrial protein import stress, in contrast to protein misfolding diseases where chronic translational repression is neurotoxic (Moreno *et al*, 2013; Radford *et al*, 2015; Celardo *et al*, 2016), and provides a mechanistic basis for the context-dependent outcomes of ISR signalling in neurodegeneration.

Import stress converts the somatic mitochondrial network from an interconnected tubular morphology into fragmented individual organelles, many of which adopt donut-shaped (toroidal) configurations visible by live imaging with the import-independent outer membrane marker mito^OMM^. This transformation is severe: import-stressed somata contain more discrete mitochondrial objects of reduced volume and increased sphericity, and with sustained expression nearly all motoneuron somata display donut-shaped mitochondria in place of the normal tubular network. These structures were not observed after chemical fixation, either with mito^OMM^ or by immunostaining for inner membrane proteins, consistent with osmotic sensitivity of the toroidal configuration (Long *et al*, 2015) and suggesting that donut-shaped mitochondria may be substantially underrepresented in studies that rely on conventional immunohistochemical analyses. Donut-shaped mitochondria have been reported in cultured non-neuronal cells under oxidative stress and hypoxia-reoxygenation, where they form through matrix swelling-induced autofusion after partial detachment from the cytoskeleton (Liu & Hajnóczky, 2011; Ahmad *et al*, 2013). *In vivo*, the only neuronal observation is in aged rhesus monkey prefrontal cortex, where mitochondrial donut frequency in presynaptic boutons inversely correlates with working memory performance (Hara *et al*, 2014); the molecular cause, however, was unknown. Our data establish a direct causal link between a defined mitochondrial protein import stress and donut formation in neurons *in vivo*. The size of these structures, which substantially exceeds that of normal axonal mitochondria, together with their resistance to PINK1/Parkin-mediated mitophagy (Zhou *et al*, 2020), provides a structural basis for their progressive somatic accumulation and the consequent depletion of mitochondria from the synaptic terminal. Whether donut formation represents an adaptive response analogous to the mitochondrial-derived compartments that sequester surplus outer membrane proteins in yeast (Wilson *et al*, 2024) remains to be determined.

The morphological transformation is accompanied by a broad transcriptional programme that intensifies progressively from Day 1 to Day 3 of mitochondrial protein import blockade. The *Drosophila Hsp70* locus provides an internal control for the temperature shift required by the TARGET system: despite encoding near-identical proteins, the 87A and 87C clusters differ in their regulatory elements (Gong & Golic, 2006; Lakhotia & Prasanth, 2002) and show divergent temporal behaviour. The 87A genes and *Hsp68* were strongly induced at Day 1 but returned to baseline by Day 3, consistent with thermal acclimation, whereas all four 87C locus genes showed sustained and progressive induction consistent with a specific role in counteracting mitochondrial protein import stress. This divergence within a single gene family validates the Day 3 transcriptome as a readout of import stress rather than residual heat shock.

This import stress-specific programme shares core features with the mitoprotein-induced stress response in yeast (Boos *et al*, 2019): sustained *Hsp70* induction, coordinate repression of nuclear-encoded OXPHOS subunits, and reduction of cytosolic ribosomal genes. That mitochondrial stress drives cytosolic chaperone remodelling across kingdoms is consistent with recent work identifying HSP70 activation through cytosolic precursor accumulation as a conserved UPR^mt^ signal in human cells (Sutandy *et al*, 2023) and HSF-1-dependent maintenance of cytosolic proteostasis during mitochondrial stress in *C. elegans* (Labbadia *et al*, 2017). The response diverges at a critical node: in yeast, *Rpn4* drives proteasome upregulation as the second wave of the import stress response, whereas *Drosophila* motoneurons, which lack an *Rpn4* orthologue, show proteasome repression. This divergence may be functionally significant. *Rpn4*-deficient yeast cells tolerate clogger stress better than wild-type by sequestering precursors into chaperone-controlled cytosolic granules rather than degrading them (Krämer *et al*, 2023). The sustained *Hsp70* induction and absent proteasome upregulation in *Drosophila* motoneurons suggest a similar sequestration-based strategy.

The transcriptional reprogramming has direct functional consequences at the synaptic terminal. Clogger-expressing NMJs show reduced quantal size, impaired evoked neurotransmitter release, and a failure to sustain transmission during prolonged stimulation. The reduction in quantal size cannot be explained by bioenergetic deficit alone, because *miro* mutant terminals that completely lack mitochondria maintain normal quantal size (Guo *et al*, 2005). Instead, multiple V-ATPase subunits are transcriptionally repressed during import stress, and V-ATPase activity is required for synaptic vesicle acidification and neurotransmitter loading at the *Drosophila* NMJ (Dulac *et al*, 2021; Hiesinger *et al*, 2005). This transcriptional repression of vesicle filling machinery provides a potential explanation for why import-stressed terminals are impaired at baseline, whereas *miro* mutant terminals fail only during sustained stimulation when ATP-dependent reserve pool mobilisation becomes limiting (Verstreken *et al*, 2005). Interestingly, neuronal depletion of Complex I in *Drosophila* motoneurons causes mitochondrial fragmentation and loss from the terminal, yet neurotransmission is maintained through a ROS-dependent homeostatic programme that enhances active zone function (Mallik *et al*, 2025). Our system instead shows repression of antioxidant defences, an expression profile inconsistent with such a compensatory mechanism.

Synaptic degeneration during import stress is not determined by local mitochondrial presence: boutons devoid of mitochondria frequently retain presynaptic active zones, and even complete absence of presynaptic mitochondria in *miro* mutants does not cause structural degeneration (Guo *et al*, 2005; Massaro *et al*, 2009). The transcriptomic upregulation of PEK/PERK during import stress implicates the integrated stress response in the degenerative process. PERK-mediated eIF2α phosphorylation attenuates global translation initiation, which would progressively deplete synaptic proteins that require continuous production and delivery to maintain synaptic integrity. Our data provide direct evidence for this link: PERK overexpression in motoneurons, in the absence of any mitochondrial perturbation, is sufficient to cause synaptic degeneration. That unrestrained ISR activation is itself harmful is independently supported by the finding that mutations in the SIFI E3 ligase, which silences DELE1-HRI signalling by targeting both proteins for degradation, cause neurodegeneration in mammalian neurons (Haakonsen *et al*, 2024). Import-stressed motoneurons transcriptionally upregulate many stability factors, including cytoskeletal organisers and kinases that oppose degeneration (Bulat *et al*, 2014; Mushtaq *et al*, 2022; Pielage *et al*, 2005; Eaton & Davis, 2005; Stephan *et al*, 2015), potentially compensating for reduced translational output by increasing transcript availability. That degeneration nonetheless occurs suggests this transcriptional compensation is insufficient to offset the translational attenuation imposed by the ISR. The endogenous transcriptional upregulation of PERK during import stress therefore places the ISR at a mechanistic crossroads: the translational attenuation that reduces precursor production simultaneously limits the synthesis of the maintenance proteins whose depletion drives synaptic disassembly.

Both GSK2606414 (GSK) and ISRIB exacerbate degeneration during import stress, while neither affects synaptic integrity in controls consistent with our RNAi-mediated PEK downregulation. Because GSK inhibits PERK directly but has off-target activity against RIPK1 (Rojas-Rivera *et al*, 2017), the convergence with ISRIB, which activates eIF2B downstream of eIF2α phosphorylation (Sidrauski *et al*, 2015), provides pharmacological evidence that the protective mechanism operates through eIF2α-mediated translational control. Restoring translation via PERK inhibition or ISRIB likely increases production of mitochondrial precursors that cannot be imported, thereby accelerating rather than relieving the degenerative process. The concurrent transcriptional upregulation of the ER stress sensor *Ire1* alongside both eIF2α kinases is consistent with ER stress activation during import stress, as the ER unfolded protein response buffers non-imported mitochondrial proteins in yeast (Knöringer *et al*, 2023). In mammalian cells, mitochondrial stress is relayed to eIF2α through the OMA1-DELE1-HRI pathway (Guo *et al*, 2020; Fessler *et al*, 2020; Anderson & Haynes, 2020), while a broader mitochondrial-to-nuclear stress response through the UPR^mt^ has been described in *C. elegans* (Haynes & Ron, 2010). *Drosophila* lacks an HRI orthologue; the activation of PEK/PERK during import stress therefore represents an alternative route to eIF2α phosphorylation, potentially enabled by the localisation of PERK at mitochondria-associated ER membranes (Verfaillie *et al*, 2012). That different kinases relay import stress to translational control in different organisms reinforces eIF2α phosphorylation itself as the conserved protective effector.

The protective role of PERK upregulation during import stress contrasts with protein misfolding and respiratory chain diseases, where PERK inhibition is beneficial. In the *Drosophila* pink1/parkin model, GSK relieves translational repression caused by aberrant ER-mitochondria contacts and is neuroprotective (Celardo *et al*, 2016). In prion-diseased mice, GSK restores synaptic protein production and prevents neurodegeneration (Moreno *et al*, 2013). In these conditions, restoring translation is beneficial because the neuron requires ongoing protein synthesis to maintain synaptic function, and translational repression does not address the upstream pathology. During import stress, by contrast, the precursor burden arises from proteins that cannot enter mitochondria, and translational attenuation directly reduces this burden. This distinction provides a principle for predicting the outcome of ISR modulation: the benefit of translational attenuation depends on whether ongoing protein synthesis contributes to the proteotoxic load or is required to counteract it. Consistent with this principle, PERK haploinsufficiency in SOD1-G85R mice, where ongoing translation directly produces the misfolding-prone protein, accelerates disease onset and increases aggregation (Wang *et al*, 2011). Several disease-associated proteins directly obstruct mitochondrial import channels (reviewed in Pfanner *et al*, 2025), but in each case import stress is accompanied by additional pathogenic mechanisms. The ANT1 clogger mouse, the most direct disease model for import channel obstruction, develops neurodegeneration in the spinal cord yet shows no ISR activation in skeletal muscle, instead engaging FOXO-dependent pathways (Coyne *et al*, 2023), indicating that the cellular response to import clogging is tissue-dependent. The convergent transcriptional reduction of eIF2B subunits alongside eIF2α kinase upregulation that characterises import-stressed motoneurons in our system has also been reported in Alzheimer’s disease brain tissue (Oliveira *et al*, 2021), raising the possibility that the translational control programme described here operates in human neurodegeneration. Our data establish that import stress represents a category of mitochondrial dysfunction for which the neuron’s endogenous defence is to translate less, not more.

## Acknowledgments

We thank the Bloomington Drosophila Stock Center (NIH P40OD018537) for fly stocks, the Developmental Studies Hybridoma Bank (DSHB, University of Iowa) for antibodies. We thank all members of the Pielage lab for helpful discussions. This work was supported by the Deutsche Forschungsgemeinschaft (DFG; GRK2737 STRESSistance) (J.M.H. and J.P.) and the Forschungsinitiative Rheinland-Pfalz BioComp (J.M.H. and J.P.).

## Author contributions

J.E. and J.P. conceived and designed the study. J.E. performed all experiments and analysed all data with assistance from M.B. L.M.L. performed electrophysiological recordings. A.G. generated transgenic constructs and performed initial experiments. S.L. performed larval locomotion assays. M.P., G.G., and M.S. performed RNA-seq library preparation, sequencing, and analysis. J.M.H. provided guidance on experimental design. J.E. and J.P. wrote the manuscript with input from all authors.

## Data availability

RNA-seq data will be deposited in the Gene Expression Omnibus (GEO) upon publication. All other data supporting the findings of this study are available from the corresponding author upon reasonable request.

## Methods

### Fly stocks

*Drosophila melanogaster* stocks used in this study are listed in Table 1. Stocks were obtained from the Bloomington *Drosophila* Stock Center (BDSC) unless otherwise indicated. All stocks were raised on standard Cold Spring Harbor Laboratory (CSHL) fly food at room temperature. Genetic crosses involving clogger constructs (cyt^DHFR^, mito^DHFR^, mito-IMM^DHFR^) were maintained at 18°C to prolong larval development and maximise the duration of construct expression. Crosses for pharmacological experiments (GSK2606414 and ISRIB) were maintained at 25°C. All other crosses were maintained at 25°C with 65 ± 5% humidity. For temporal control of transgene expression, the TARGET system (McGuire *et al*, 2003) was used: animals carrying tub-Gal80^ts^ were raised at 18°C and shifted to 29°C for the indicated duration prior to analysis at the wandering third instar larval stage. For control genotypes, *w*^1118^ was crossed to the respective Gal4 driver unless otherwise indicated.

**Table 1:**
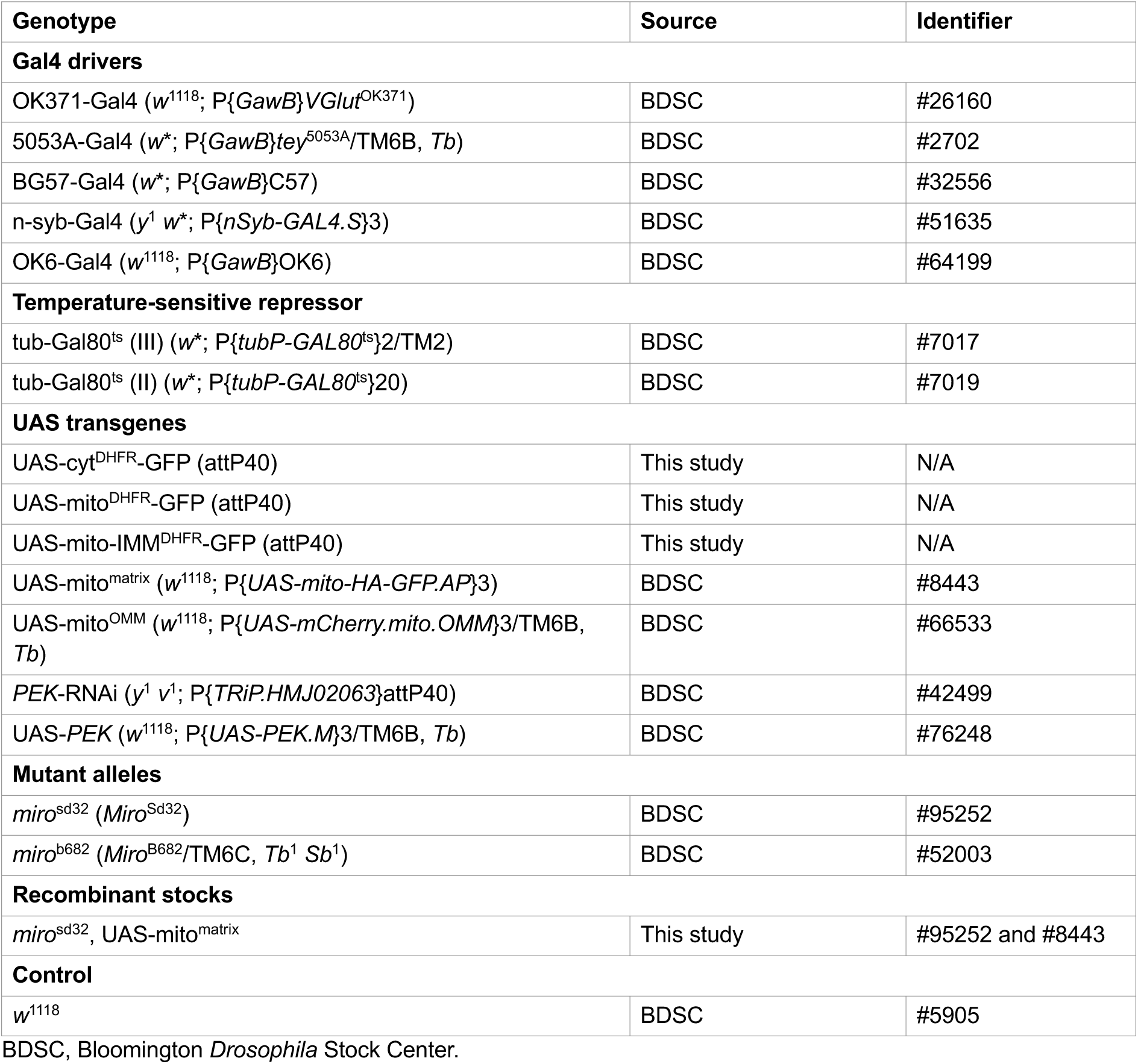
*Drosophila* stocks used in this study.

### Generation of clogger constructs

Clogger constructs were adapted from the yeast import stress system (Boos et al., 2019). The coding sequences for cyt-DHFR, b2Δ-DHFR (mito^DHFR^), and b2-DHFR (mito-IMM^DHFR^) were amplified by PCR from the yeast expression plasmids pYX233-cytDHFR, pYX233-b2Δ-DHFR, and pYX233-b2-DHFR using primers flanked by attB recombination sites. PCR products were cloned into pDONR221 via Gateway BP reaction to generate entry clones. Expression clones were produced by LR recombination with the destination vector pUAST-10xUAS-rfA-EGFP, resulting in constructs carrying 10 tandem UAS repeats upstream of the respective DHFR coding sequence fused to a C-terminal GFP tag. cyt^DHFR^ lacks any mitochondrial targeting sequence and remains cytosolic. mito^DHFR^ carries the first 167 amino acids of yeast cytochrome b2, including the mitochondrial targeting sequence but lacking the transmembrane domain, fused to DHFR. mito-IMM^DHFR^ carries the same region of cytochrome b2 including the transmembrane domain, which serves as a stop-transfer signal at the inner mitochondrial membrane. All constructs were sequence-verified and transgenic flies were generated by phiC31-mediated transgenesis at the attP40 landing site (Bischof *et al*, 2007) to ensure equivalent expression levels. The following primers were used for amplification: cytDHFR forward 5′-CACCATGGTTCGACCATTGAACTG-3′, b2-DHFR forward 5′-CACCATGCTAAAATACAAACC-3′ (shared by mito^DHFR^ and mito-IMM^DHFR^), and DHFR reverse 5′-GTCTTTCTTCTCGTAGACTTC-3′ (shared by all three constructs); the 5′ CACC overhang enables directional Gateway cloning.

### Solutions

Dissecting saline contained 70 mM NaCl, 5 mM KCl, 10 mM MgCl_2_, 5 mM NaHCO_3_, 115 mM sucrose, 5 mM trehalose, 5 mM HEPES, and 2 mM EGTA, pH 7.0. Recording HL3 saline had the same composition without EGTA, supplemented with CaCl_2_ at the concentrations indicated for each experiment.

### Immunohistochemistry

Wandering third instar larvae were dissected in ice-cold dissecting saline and fixed in Bouin’s fixative (Carl Roth, #6482.3) for 5 minutes at room temperature. Preparations were washed in PBST (PBS containing 0.1% Triton X-100) until residual fixative was removed. Primary antibodies were applied in PBST overnight at 4°C. After five washes in PBST (10 minutes each), secondary antibodies and HRP conjugates were applied in PBST for 2 hours at room temperature in the dark, followed by five washes in PBST. Preparations were incubated in 70% glycerol (in PBS) for at least 1 hour before mounting on glass slides in Fluoromount-G (Invitrogen, #00-4958-02). The following primary antibodies were used: mouse anti-GFP (3E6) 1:2,000 (Invitrogen, #A-11120), mouse anti-Brp (nc82) 1:250 (Developmental Studies Hybridoma Bank [DSHB]), mouse anti-HA 1:500 (Roche, #11583816001), and rabbit anti-GluRIIc 1:3,000 (Pielage *et al*, 2011). The following secondary antibodies and conjugates were used: goat anti-mouse Alexa Fluor 488 1:1,000 (Invitrogen, #A11029), goat anti-rabbit Alexa Fluor 568 1:1,000 (Invitrogen, #A-11036), goat anti-HRP Cy3 1:1,000 (Jackson ImmunoResearch, #123-165-021), and goat anti-HRP Alexa Fluor 647 1:500 (Jackson ImmunoResearch, #123-605-021). All genotypes within a single experiment were stained together and imaged under identical acquisition settings.

### Live imaging

Wandering third instar larvae were dissected in ice-cold dissecting saline on a small piece of Sylgard. Preparations were rinsed twice in dissecting saline and excess liquid was removed. The Sylgard block was inverted onto a 35 mm glass-bottom dish so that the opened preparation was pressed against the glass. Images were acquired on a Stellaris 8 confocal microscope (Leica) using a 63×/1.4 NA oil immersion objective. Z-stacks were processed with the Leica Lightning deconvolution software using standard adaptive settings. For axonal live imaging of mitochondrial transport, recordings were acquired for 2 minutes at 1 frame per second with Leica dynamic signal enhancement (Frames = 11, Weight = 4) followed by Lightning deconvolution. Kymographs were generated using the Multi Kymograph function in Fiji (ImageJ; Schindelin et al., 2012).

### MitoTracker staining

Wandering third instar larvae expressing clogger constructs were dissected as for live imaging. Before mounting, preparations were incubated in dissecting saline containing 500 nM MitoTracker Red CMXRos (Invitrogen, #M7512) and goat anti-HRP Alexa Fluor 647 (1:500, Jackson ImmunoResearch) for 5 minutes. After rinsing once in dissecting saline, preparations were mounted on glass-bottom dishes and imaged live at muscle 6/7 NMJs to capture MitoTracker, GFP-tagged construct, and neuronal membrane signals. Preparations were subsequently fixed in Bouin’s fixative for 5 minutes and processed for immunohistochemistry against Brp and GluRIIc as described above. The same NMJs were re-imaged after antibody staining, and live and fixed images were aligned using the HRP signal with the Landmark Correspondences function in Fiji. The neuronal membrane signal was used to distinguish presynaptic from postsynaptic (muscle) mitochondria.

### Electrophysiological recordings

Intracellular recordings were performed on wandering third instar larvae dissected in ice-cold dissecting saline (see Solutions). Single-electrode current-clamp recordings were performed with sharp electrodes filled with 1 M KCl (resistance 15–20 MΩ) on muscle 6 of abdominal segments A3 and A4 in recording HL3 containing 0.5 mM CaCl_2_ for baseline measurements or 1.0 mM CaCl_2_ for high-frequency stimulation. Miniature excitatory junction potentials (mEJPs) were recorded for 60 seconds. Evoked excitatory junction potentials (EJPs) were elicited by stimulating the segmental nerve with 30 suprathreshold pulses (6 V, 3 ms duration) at 1 Hz, delivered by an isolated pulse stimulator (Model 2100, A-M Systems). Quantal content was estimated by dividing the mean EJP amplitude by the mean mEJP amplitude. Only recordings with a resting membrane potential between −55 and −75 mV and an input resistance ≥ 3 MΩ were included. For high-frequency stimulation, NMJs were stimulated at 10 Hz for 300 seconds. Recordings were acquired using an Axoclamp 900A amplifier and digitised with a Digidata 1440A (Molecular Devices). Traces were analysed using Clampfit 11.2 (Molecular Devices).

### Larval locomotion

Larval locomotion was assessed using the frustrated total internal reflection imaging method FIM (Risse *et al*, 2013) and analysed with FIMtrack v3.1.32.5 software (Risse *et al*, 2014). Wandering third instar larvae were collected and rested on a 1.5% agarose plate for 2 minutes at room temperature. Five animals at a time were placed in the centre of a 1.5% agarose plate on a FIM table. Larval motion was recorded in the dark at room temperature for 2 minutes at 10 Hz using a Basler acA2040-90um camera and Basler Pylon v7.2.1 software. Ten rounds of five animals each were recorded per genotype in a single session. Recorded tracks were analysed and quantified using FIMtrack. Each data point in the quantifications represents the average of five animals from a single run.

### Pharmacological treatment

For pharmacological inhibition of PERK, standard fly food was heated until liquefied and supplemented with GSK2606414 (Merck) dissolved in DMSO to a final concentration of 10 µM. For ISRIB treatment, food was supplemented with ISRIB (Merck, #SML0843) dissolved in DMSO to a final concentration of 5 nM. Approximately 5 ml portions were distributed into fresh food vials. Vehicle control food was prepared identically using equivalent volumes of DMSO. Parental crosses were placed on drug-containing or control food at 25°C, and the resulting progeny were exposed to the compounds throughout development until dissection at the wandering third instar larval stage.

### Quantification of synaptic degeneration

Synaptic degeneration was quantified using an established assay (Eaton *et al*, 2002; Pielage *et al*, 2005; Mushtaq *et al*, 2022). NMJs were stained for the presynaptic active zone marker Brp and the postsynaptic glutamate receptor GluRIIc. Boutons retaining a postsynaptic GluRIIc signal without a corresponding presynaptic Brp signal were scored as degenerated. Quantification was performed on muscle 4 and muscle 6/7 NMJs of hemisegments A2–A6 using a TCS SPE DM5500Q microscope (Leica) with a 40×/0.75 NA air objective. The frequency of synaptic degeneration represents the percentage of NMJs per animal displaying at least one degenerated bouton. Degeneration severity is reported as the number of degenerated boutons per NMJ. NMJ elimination was defined as near complete loss of Brp with complete fragmentation of the neuronal membrane. Representative images were acquired on a Stellaris 8 confocal microscope (Leica) with a 63×/1.4 NA oil immersion objective and processed with Lightning deconvolution.

### Quantification of mitochondrial parameters in the soma

For analysis of mito^matrix^-labelled mitochondria in the soma, mito^matrix^ was co-expressed with clogger constructs in motoneurons, and wandering third instar larvae were dissected and stained with anti-HA antibodies. Images were acquired on a Stellaris 8 confocal microscope with a 63×/1.4 NA oil immersion objective and processed with Lightning deconvolution. Individual somata were isolated by three-dimensional cropping in the Leica LAS X 3D Visualisation tool. Mitochondrial parameters (number, volume, surface area, sphericity, and signal intensity) were measured using the MitochondriaAnalyzer plugin for Fiji (Chaudhry *et al*, 2020) with standard thresholding settings (Block Size 1.45 µm, C-Value 9). Signal intensities were measured by applying the threshold masks to the original, non-deconvolved confocal images.

For analysis of mito^OMM^-labelled mitochondria, wandering third instar larvae co-expressing mito^OMM^ and mito-IMM^DHFR^ were dissected and imaged live as described above. Individual somata were isolated as above and mitochondrial parameters were measured using MitochondriaAnalyzer (Block Size 1.5 µm, C-Value 11). Donut-shaped mitochondria were counted manually in the mito^OMM^channel. For analysis of early stages of import stress, expression was induced via the TARGET system by shifting third instar larvae from 18°C to 29°C for 15 hours, and donut structures were counted manually.

### Quantification of mitochondrial parameters at the NMJ

Images of muscle 4 NMJs from hemisegments A3–A5 were acquired on a Stellaris 8 confocal microscope with a 63×/1.4 NA oil immersion objective under identical settings across genotypes. NMJ area was defined by the neuronal membrane (HRP) signal using the default auto-threshold function in Fiji. Mitochondrial signals within the NMJ were thresholded with manual adjustment to account for genotype-dependent differences in mito^matrix^ signal intensity. Mitochondrial density, size, and signal intensity were measured using the standard Measure function in Fiji. All thresholds and masks were manually validated.

### Transcriptomic analysis

For RNA-seq, mito-IMM^DHFR^ expression was induced pan-neuronally using the TARGET system (n-syb-Gal4; tub-Gal80^ts^) by shifting animals from 18°C to 29°C for 1, 2, or 3 days prior to collection at the wandering third instar larval stage. Uninduced controls (0 days) were maintained at 18°C. For each condition, 15 brains from wandering third instar larvae were dissected and pooled per biological replicate from 3 independent crosses. Brains were transferred into 1.5 ml tubes containing 250 µl peqGOLD TriFast (peqlab) and homogenised. RNA was isolated according to manufacturer’s instructions, digested with DNase with subsequent purification with acid phenol. After integrity check with the Bioanalyzer RNA Pico Chips (Agilent), Illumina-compatible RNA libraries were prepared using the NEBNext Ultra II Directional RNA Library Prep Kit for Illumina (New England Biolabs) with the NEBNext Poly(A) mRNA Magnetic Isolation Module (New England Biolabs). Sequencing was performed on a HiSeq 2500 (Illumina) with single-end 50 bp reads. Adapter trimming was performed with Trim Galore (default settings). Reads were mapped to the *Drosophila melanogaster* genome using Bowtie2 (Langmead & Salzberg, 2012) (end-to-end, high-sensitivity mode). Expression levels were calculated using the Geneious “calculate expression” function; non-coding RNA and pseudogenes were excluded. Differential expression analysis was performed using the DESeq2 plugin (Love *et al*, 2014) in Geneious (fit-type: parametric; significance thresholds: adjusted p-value < 0.01, |log_2_ fold change| > 1). Gene ontology terms were extracted from the PANTHER classification system. Volcano plots were generated in R (v4.2.2) using the ggplot2 package (v3.5.2) (Wickham, 2016). n = 2 biological replicates for 0 days, n = 3 biological replicates for 1–3 days.

### Microscopy and image processing

Confocal images for all figures were acquired on a Stellaris 8 confocal microscope (Leica) with a 63×/1.4 NA oil immersion objective unless otherwise stated. Screening and quantification of synaptic degeneration were performed on a TCS SPE DM5500Q microscope (Leica) with a 40×/0.75 NA air objective. Images were processed in Fiji (ImageJ) (Schindelin *et al*, 2012) by cropping, rotation, and linear adjustments of brightness. Figures and schematics were created in Affinity Designer 2 (v2.6.3, Serif Europe Ltd.).

### Statistical analysis

Statistical analyses were performed using GraphPad Prism 10. All data were tested for normal distribution using the Shapiro-Wilk test. For normally distributed data, one-way ANOVA with Dunnett’s post-hoc correction was applied. For non-normally distributed data, Kruskal-Wallis tests with Dunn’s post-hoc correction were used. Two-group comparisons were performed using unpaired t-tests or Mann-Whitney tests as appropriate. All tests were two-tailed. Significance levels were defined as: ***p ≤ 0.001, **p ≤ 0.01, *p ≤ 0.05, and ns p ≥ 0.05.

## References

1. Ahmad T, Aggarwal K, Pattnaik B, Mukherjee S, Sethi T, Tiwari BK, Kumar M, Micheal A, Mabalirajan U, Ghosh B, et al (2013) Computational classification of mitochondrial shapes reflects stress and redox state. Cell Death Dis 4: e461

2. Anderson DD, Woeller CF, Chiang E-P, Shane B & Stover PJ (2012) Serine hydroxymethyltransferase anchors de novo thymidylate synthesis pathway to nuclear lamina for DNA synthesis. J Biol Chem 287: 7051–7062

3. Anderson NS & Haynes CM (2020) Folding the mitochondrial UPR into the integrated stress response. Trends Cell Biol 30: 428–439

4. Bischof J, Maeda RK, Hediger M, Karch F & Basler K (2007) An optimized transgenesis system for Drosophila using germ-line-specific phiC31 integrases. Proc Natl Acad Sci U S A 104: 3312–3317

5. Boos F, Krämer L, Groh C, Jung F, Haberkant P, Stein F, Wollweber F, Gackstatter A, Zöller E, van der Laan M, et al (2019) Mitochondrial protein-induced stress triggers a global adaptive transcriptional programme. Nat Cell Biol 21: 442–451

6. Bulat V, Rast M & Pielage J (2014) Presynaptic CK2 promotes synapse organization and stability by targeting Ankyrin2. J Cell Biol 204: 77–94

7. Celardo I, Costa AC, Lehmann S, Jones C, Wood N, Mencacci NE, Mallucci GR, Loh SHY & Martins LM (2016) Mitofusin-mediated ER stress triggers neurodegeneration in pink1/parkin models of Parkinson’s disease. Cell Death Dis 7: e2271

8. Cenini G, Rüb C, Bruderek M & Voos W (2016) Amyloid β-peptides interfere with mitochondrial preprotein import competence by a coaggregation process. Mol Biol Cell 27: 3257–3272

9. Chaudhry A, Shi R & Luciani DS (2020) A pipeline for multidimensional confocal analysis of mitochondrial morphology, function, and dynamics in pancreatic β-cells. Am J Physiol Endocrinol Metab 318: E87–E101

10. Coyne LP, Wang X, Song J, de Jong E, Schneider K, Massa PT, Middleton FA, Becker T & Chen XJ (2023) Mitochondrial protein import clogging as a mechanism of disease. Elife 12

11. Desai S, Grefte S, van de Westerlo E, Lauwen S, Paters A, Prehn JHM, Gan Z, Keijer J, Adjobo-Hermans MJW & Koopman WJH (2024) Performance of TMRM and Mitotrackers in mitochondrial morphofunctional analysis of primary human skin fibroblasts. Biochim Biophys Acta Bioenerg 1865: 149027

12. Devi L, Prabhu BM, Galati DF, Avadhani NG & Anandatheerthavarada HK (2006) Accumulation of amyloid precursor protein in the mitochondrial import channels of human Alzheimer’s disease brain is associated with mitochondrial dysfunction. J Neurosci 26: 9057–9068

13. Di Maio R, Barrett PJ, Hoffman EK, Barrett CW, Zharikov A, Borah A, Hu X, McCoy J, Chu CT, Burton EA, et al (2016) α-Synuclein binds to TOM20 and inhibits mitochondrial protein import in Parkinson’s disease. Sci Transl Med 8: 342ra78

14. Dulac A, Issa A-R, Sun J, Matassi G, Jonas C, Chérif-Zahar B, Cattaert D & Birman S (2021) A novel neuron-specific regulator of the V-ATPase in Drosophila. eNeuro 8: ENEURO.0193-21.2021

15. Dutta D, Briere LC, Kanca O, Marcogliese PC, Walker MA, High FA, Vanderver A, Krier J, Carmichael N, Callahan C, et al (2020) De novo mutations in TOMM70, a receptor of the mitochondrial import translocase, cause neurological impairment. Hum Mol Genet 29: 1568–1579

16. Eaton BA & Davis GW (2005) LIM Kinase1 controls synaptic stability downstream of the type II BMP receptor. Neuron 47: 695–708

17. Eaton BA, Fetter RD & Davis GW (2002) Dynactin is necessary for synapse stabilization. Neuron 34: 729–741

18. Enneking E-M, Kudumala SR, Moreno E, Stephan R, Boerner J, Godenschwege TA & Pielage J (2013) Transsynaptic coordination of synaptic growth, function, and stability by the L1-type CAM Neuroglian. PLoS Biol 11: e1001537

19. Fessler E, Eckl E-M, Schmitt S, Mancilla IA, Meyer-Bender MF, Hanf M, Philippou-Massier J, Krebs S, Zischka H & Jae LT (2020) A pathway coordinated by DELE1 relays mitochondrial stress to the cytosol. Nature 579: 433–437

20. Forgac M (2007) Vacuolar ATPases: rotary proton pumps in physiology and pathophysiology. Nat Rev Mol Cell Biol 8: 917–929

21. Gong WJ & Golic KG (2006) Loss of Hsp70 in Drosophila is pleiotropic, with effects on thermotolerance, recovery from heat shock and neurodegeneration. Genetics 172: 275–286

22. Graf ER, Heerssen HM, Wright CM, Davis GW & DiAntonio A (2011) Stathmin is required for stability of the Drosophila neuromuscular junction. J Neurosci 31: 15026–15034

23. Guo X, Aviles G, Liu Y, Tian R, Unger BA, Lin Y-HT, Wiita AP, Xu K, Correia MA & Kampmann M (2020) Mitochondrial stress is relayed to the cytosol by an OMA1-DELE1-HRI pathway. Nature 579: 427–432

24. Guo X, Macleod GT, Wellington A, Hu F, Panchumarthi S, Schoenfield M, Marin L, Charlton MP, Atwood HL & Zinsmaier KE (2005) The GTPase dMiro is required for axonal transport of mitochondria to Drosophila synapses. Neuron 47: 379–393

25. Haakonsen DL, Heider M, Ingersoll AJ, Vodehnal K, Witus SR, Uenaka T, Wernig M & Rapé M (2024) Stress response silencing by an E3 ligase mutated in neurodegeneration. Nature 626: 874–880

26. Hara Y, Yuk F, Puri R, Janssen WGM, Rapp PR & Morrison JH (2014) Presynaptic mitochondrial morphology in monkey prefrontal cortex correlates with working memory and is improved with estrogen treatment. Proc Natl Acad Sci U S A 111: 486–491

27. Haynes CM & Ron D (2010) The mitochondrial UPR – protecting organelle protein homeostasis. J Cell Sci 123: 3849–3855

28. Hetz C & Mollereau B (2014) Disturbance of endoplasmic reticulum proteostasis in neurodegenerative diseases. Nat Rev Neurosci 15: 233–249

29. Hiesinger PR, Fayyazuddin A, Mehta SQ, Rosenmund T, Schulze KL, Zhai RG, Verstreken P, Cao Y, Zhou Y, Kunz J, et al (2005) The v-ATPase V0 subunit a1 is required for a late step in synaptic vesicle exocytosis in Drosophila. Cell 121: 607–620

30. Hurd DD & Saxton WM (1996) Kinesin mutations cause motor neuron disease phenotypes by disrupting fast axonal transport in Drosophila. Genetics 144: 1075–1085

31. Knöringer K, Groh C, Krämer L, Stein KC, Hansen KG, Zimmermann J, Morgan B, Herrmann JM, Frydman J & Boos F (2023) The unfolded protein response of the endoplasmic reticulum supports mitochondrial biogenesis by buffering nonimported proteins. Mol Biol Cell 34: ar95

32. Koch I, Schwarz H, Beuchle D, Goellner B, Langegger M & Aberle H (2008) Drosophila ankyrin 2 is required for synaptic stability. Neuron 58: 210–222

33. Koehler CM, Leuenberger D, Merchant S, Renold A, Junne T & Schatz G (1999) Human deafness dystonia syndrome is a mitochondrial disease. Proc Natl Acad Sci U S A 96: 2141–2146

34. Krämer L, Dalheimer N, Räschle M, Storchová Z, Pielage J, Boos F & Herrmann JM (2023) MitoStores: chaperone-controlled protein granules store mitochondrial precursors in the cytosol. EMBO J 42: e112309

35. Labbadia J, Brielmann RM, Neto MF, Lin Y-F, Haynes CM & Morimoto RI (2017) Mitochondrial stress restores the heat shock response and prevents proteostasis collapse during aging. Cell Rep 21: 1481–1494

36. Lakhotia SC & Prasanth KV (2002) Tissue– and development-specific induction and turnover of hsp70 transcripts from loci 87A and 87C after heat shock and during recovery in Drosophila melanogaster. J Exp Biol 205: 345–358

37. Langmead B & Salzberg SL (2012) Fast gapped-read alignment with Bowtie 2. Nat Methods 9: 357–359

38. Liu X & Hajnóczky G (2011) Altered fusion dynamics underlie unique morphological changes in mitochondria during hypoxia-reoxygenation stress. Cell Death Differ 18: 1561–1572

39. Long Q, Zhao D, Fan W, Yang L, Zhou Y, Qi J, Wang X & Liu X (2015) Modeling of mitochondrial donut formation. Biophys J 109: 892–899

40. Love MI, Huber W & Anders S (2014) Moderated estimation of fold change and dispersion for RNA-seq data with DESeq2. Genome Biol 15: 550

41. Mallik B, Bhat SA, Wang X & Frank CA (2025) Mitochondrial Complex I and ROS control neuromuscular function through opposing pre– and postsynaptic mechanisms. PLoS Biol 23: e3003388

42. Malzer E, Daly M-L, Moloney A, Sendall TJ, Thomas SE, Ryder E, Ryoo HD, Crowther DC, Lomas DA & Marciniak SJ (2010) Impaired tissue growth is mediated by checkpoint kinase 1 (CHK1) in the integrated stress response. J Cell Sci 123: 2892–2900

43. Mårtensson CU, Priesnitz C, Song J, Ellenrieder L, Doan KN, Boos F, Floerchinger A, Zufall N, Oeljeklaus S, Warscheid B, et al (2019) Mitochondrial protein translocation-associated degradation. Nature 569: 679–683

44. Massaro CM, Pielage J & Davis GW (2009) Molecular mechanisms that enhance synapse stability despite persistent disruption of the spectrin/ankyrin/microtubule cytoskeleton. J Cell Biol 187: 101–117

45. McGuire SE, Le PT, Osborn AJ, Matsumoto K & Davis RL (2003) Spatiotemporal rescue of memory dysfunction in Drosophila. Science 302: 1765–1768

46. Moreno JA, Halliday M, Molloy C, Radford H, Verity N, Axten JM, Ortori CA, Willis AE, Fischer PM, Barrett DA, et al (2013) Oral treatment targeting the unfolded protein response prevents neurodegeneration and clinical disease in prion-infected mice. Sci Transl Med 5: 206ra138

47. Mushtaq Z, Aavula K, Lasser DA, Kieweg ID, Lion LM, Kins S & Pielage J (2022) Madm/NRBP1 mediates synaptic maintenance and neurodegeneration-induced presynaptic homeostatic potentiation. Cell Rep 41: 111710

48. Nowicka U, Chroscicki P, Stroobants K, Sladowska M, Turek M, Uszczynska-Ratajczak B, Kundra R, Goral T, Perni M, Dobson CM, et al (2021) Cytosolic aggregation of mitochondrial proteins disrupts cellular homeostasis by stimulating the aggregation of other proteins. Elife 10

49. Oliveira MM, Lourenco MV, Longo F, Kasica NP, Yang W, Ureta G, Ferreira DDP, Mendonça PHJ, Bernales S, Ma T, et al (2021) Correction of eIF2-dependent defects in brain protein synthesis, synaptic plasticity, and memory in mouse models of Alzheimer’s disease. Sci Signal 14: eabc5429

50. Pfanner N, den Brave F & Becker T (2025) Mitochondrial protein import stress. Nat Cell Biol 27: 188–201

51. Pielage J, Bulat V, Zuchero JB, Fetter RD & Davis GW (2011) Hts/Adducin controls synaptic elaboration and elimination. Neuron 69: 1114–1131

52. Pielage J, Cheng L, Fetter RD, Carlton PM, Sedat JW & Davis GW (2008) A presynaptic giant ankyrin stabilizes the NMJ through regulation of presynaptic microtubules and transsynaptic cell adhesion. Neuron 58: 195–209

53. Pielage J, Fetter RD & Davis GW (2005) Presynaptic spectrin is essential for synapse stabilization. Curr Biol 15: 918–928

54. Pilling AD, Horiuchi D, Lively CM & Saxton WM (2006) Kinesin-1 and Dynein are the primary motors for fast transport of mitochondria in Drosophila motor axons. Mol Biol Cell 17: 2057–2068

55. Radford H, Moreno JA, Verity N, Halliday M & Mallucci GR (2015) PERK inhibition prevents tau-mediated neurodegeneration in a mouse model of frontotemporal dementia. Acta Neuropathol 130: 633–642

56. Rangaraju V, Calloway N & Ryan TA (2014) Activity-driven local ATP synthesis is required for synaptic function. Cell 156: 825–835

57. Risse B, Otto N, Berh D, Jiang X & Klämbt C (2014) FIM imaging and FIMtrack: two new tools allowing high-throughput and cost effective locomotion analysis. J Vis Exp

58. Risse B, Thomas S, Otto N, Löpmeier T, Valkov D, Jiang X & Klämbt C (2013) FIM, a novel FTIR-based imaging method for high throughput locomotion analysis. PLoS One 8: e53963

59. Rojas-Rivera D, Delvaeye T, Roelandt R, Nerinckx W, Augustyns K, Vandenabeele P & Bertrand MJM (2017) When PERK inhibitors turn out to be new potent RIPK1 inhibitors: critical issues on the specificity and use of GSK2606414 and GSK2656157. Cell Death Differ 24: 1100–1110

60. Russo GJ, Louie K, Wellington A, Macleod GT, Hu F, Panchumarthi S & Zinsmaier KE (2009) Drosophila Miro is required for both anterograde and retrograde axonal mitochondrial transport. J Neurosci 29: 5443–5455

61. Schindelin J, Arganda-Carreras I, Frise E, Kaynig V, Longair M, Pietzsch T, Preibisch S, Rueden C, Saalfeld S, Schmid B, et al (2012) Fiji: an open-source platform for biological-image analysis. Nat Methods 9: 676–682

62. Sidrauski C, Tsai JC, Kampmann M, Hearn BR, Vedantham P, Jaishankar P, Sokabe M, Mendez AS, Newton BW, Tang EL, et al (2015) Pharmacological dimerization and activation of the exchange factor eIF2B antagonizes the integrated stress response. Elife 4: e07314

63. Stephan R, Goellner B, Moreno E, Frank CA, Hugenschmidt T, Genoud C, Aberle H & Pielage J (2015) Hierarchical microtubule organization controls axon caliber and transport and determines synaptic structure and stability. Dev Cell 33: 5–21

64. Sutandy FXR, Gößner I, Tascher G & Münch C (2023) A cytosolic surveillance mechanism activates the mitochondrial UPR. Nature 618: 849–854

65. Tsuyama T, Tsubouchi A, Usui T, Imamura H & Uemura T (2017) Mitochondrial dysfunction induces dendritic loss via eIF2α phosphorylation. J Cell Biol 216: 815–834

66. Vagnoni A & Bullock SL (2016) A simple method for imaging axonal transport in aging neurons using the adult Drosophila wing. Nat Protoc 11: 1711–1723

67. Verfaillie T, Rubio N, Garg AD, Bultynck G, Rizzuto R, Decuypere J-P, Piette J, Linehan C, Gupta S, Samali A, et al (2012) PERK is required at the ER-mitochondrial contact sites to convey apoptosis after ROS-based ER stress. Cell Death Differ 19: 1880–1891

68. Verstreken P, Ly CV, Venken KJT, Koh T-W, Zhou Y & Bellen HJ (2005) Synaptic mitochondria are critical for mobilization of reserve pool vesicles at Drosophila neuromuscular junctions. Neuron 47: 365–378

69. Wang L, Popko B & Roos RP (2011) The unfolded protein response in familial amyotrophic lateral sclerosis. Hum Mol Genet 20: 1008–1015

70. Wang X & Chen XJ (2015) A cytosolic network suppressing mitochondria-mediated proteostatic stress and cell death. Nature 524: 481–484

71. Wei X, Du M, Xie J, Luo T, Zhou Y, Zhang K, Li J, Chen D, Xu P, Jia M, et al (2020) Mutations in TOMM70 lead to multi-OXPHOS deficiencies and cause severe anemia, lactic acidosis, and developmental delay. J Hum Genet 65: 231–240

72. Weidberg H & Amon A (2018) MitoCPR-A surveillance pathway that protects mitochondria in response to protein import stress. Science 360: eaan4146

73. Wickham H (2016) Ggplot2: Elegant graphics for data analysis 2nd ed. Cham, Switzerland: Springer International Publishing

74. Wilson ZN, Balasubramaniam SS, Wong S, Schuler M-H, Wopat MJ & Hughes AL (2024) Mitochondrial-derived compartments remove surplus proteins from the outer mitochondrial membrane. J Cell Biol 223

75. Woeller CF, Anderson DD, Szebenyi DME & Stover PJ (2007) Evidence for small ubiquitin-like modifier-dependent nuclear import of the thymidylate biosynthesis pathway. J Biol Chem 282: 17623–17631

76. Wrobel L, Topf U, Bragoszewski P, Wiese S, Sztolsztener ME, Oeljeklaus S, Varabyova A, Lirski M, Chroscicki P, Mroczek S, et al (2015) Mistargeted mitochondrial proteins activate a proteostatic response in the cytosol. Nature 524: 485–488

77. Yablonska S, Ganesan V, Ferrando LM, Kim J, Pyzel A, Baranova OV, Khattar NK, Larkin TM, Baranov SV, Chen N, et al (2019) Mutant huntingtin disrupts mitochondrial proteostasis by interacting with TIM23. Proc Natl Acad Sci U S A 116: 16593–16602

78. Yano H, Baranov SV, Baranova OV, Kim J, Pan Y, Yablonska S, Carlisle DL, Ferrante RJ, Kim AH & Friedlander RM (2014) Inhibition of mitochondrial protein import by mutant huntingtin. Nat Neurosci 17: 822–831

79. Zhou Y, Long Q, Wu H, Li W, Qi J, Wu Y, Xiang G, Tang H, Yang L, Chen K, et al (2020) Topology-dependent, bifurcated mitochondrial quality control under starvation. Autophagy 16: 562–574

