## Supplemental Data for "Mitochondrial protein import stress causes progressive neurodegeneration opposed by PERK - eIF2α signalling"

#### Supplemental Figure Legends

##### Supplemental Figure 1: Early co-localisation of mito-IMM$^{DHFR}$ with mitochondria.

**(A)**, **(B)** Live imaging of motoneuron somata after 15 hours of mito-IMM$^{DHFR}$ (green) and mito$^{OMM}$ (magenta) co-expression using the TARGET system (tub-Gal80$^{ts}$; OK371-Gal4). **(A)** Example of a soma with a largely intact mitochondrial network. mito-IMM$^{DHFR}$ co-localises with mito$^{OMM}$ throughout the network (arrowheads). First donut-shaped structures are beginning to form (arrows). **(B)** Example of a soma with more advanced donut formation (arrows). mito-IMM$^{DHFR}$ co-localises with all mito$^{OMM}$-labelled structures at this stage. Dashed boxes indicate regions shown at higher magnification. Scale bars: 2 µm. **(C)** Live imaging of a motoneuron axon after 48 hours of mito-IMM$^{DHFR}$ (green) and mito$^{OMM}$ (magenta) co-expression. mito-IMM$^{DHFR}$ co-localises with mito$^{OMM}$ in discrete axonal mitochondria (arrowheads), confirming mitochondrial targeting in the axonal compartment. Dashed boxes indicate regions shown at higher magnification. Scale bars: 5 µm.

##### Supplemental Figure 2: Synaptic degeneration at muscle 6/7 NMJs.

**(A)**–**(D)** Confocal images of muscle 6/7 NMJs. Constructs were expressed in motoneurons (OK371-Gal4; **A**–**C**) or in somatic muscles (BG57-Gal4; **D**). Top row: NMJ overview with neuronal membrane (HRP, white). Lower rows at higher magnification: neuronal membrane (magenta), Brp (green) and GluRIIc (magenta) merge, and Brp alone. **(A)**, **(B)** cyt$^{DHFR}$ and mito$^{DHFR}$ expressing motoneurons display intact neuronal membranes with Brp and GluRIIc in precise apposition at all boutons (arrowheads). **(C)** Motoneuron expression of mito-IMM$^{DHFR}$ causes fragmentation of the neuronal membrane and loss of Brp from individual boutons despite persistent GluRIIc (arrows). **(D)** Muscle expression of mito-IMM$^{DHFR}$ does not cause synaptic degeneration. Asterisks indicate regions shown at higher magnification. Scale bars: 10 µm (overviews), 2 µm (zoom-ins). **(E)** Degeneration severity at muscle 6/7 NMJs for neuronal and muscle expression of the indicated constructs. **(F)** Degeneration severity at muscle 4 NMJs. Only neuronal expression of mito-IMM$^{DHFR}$ significantly increases degeneration severity at both muscle types. Boxplots indicate median and interquartile range. Data were tested for normality (Shapiro-Wilk) and analysed with parametric or non-parametric tests as appropriate, with post-hoc correction for multiple comparisons. ***$p \leq 0.001$. n = 160 NMJs from 16 animals (neuronal expression) or 100 NMJs from 10 animals (muscle expression) (E, F).

##### Supplemental Figure 3: Raw electrophysiological values at muscle 6/7 NMJs.

**(A)**–**(D)** Non-normalised electrophysiological recordings at muscle 6/7 NMJs of motoneurons expressing cyt$^{DHFR}$, mito$^{DHFR}$, or mito-IMM$^{DHFR}$ (OK371-Gal4). **(A)** mEJP amplitude (mV). **(B)** EJP amplitude (mV). **(C)** Quantal content (EJP/mEJP). **(D)** mEJP frequency (Hz). In mito-IMM$^{DHFR}$ expressing animals, all measured parameters are significantly reduced. mito$^{DHFR}$ does not differ from cyt$^{DHFR}$ controls. Boxplots indicate median and interquartile range; dashed grey lines represent mean control values. Data were tested for normality (Shapiro-Wilk) and analysed with parametric or non-parametric tests as appropriate, with post-hoc correction for multiple comparisons. *$p \leq 0.05$, ***$p \leq 0.001$. n = 28 (cyt$^{DHFR}$), 31 (mito$^{DHFR}$), or 38 (mito-IMM$^{DHFR}$) recordings from 10 (cyt$^{DHFR}$, mito$^{DHFR}$) or 12 (mito-IMM$^{DHFR}$) animals.

**Supplemental Figure 4: Extended transcriptomic analyses.**

**(A)** Temporal expression profiles of *Hsp70* gene family members across 0 to 3 days of mito-IMM$^{DHFR}$ expression. *Hsp70* 87A locus genes (*Hsp70Aa*, *Hsp70Ab*) and *Hsp68* are strongly induced after 1 day but return to baseline by 3 days, consistent with thermal acclimation. All four 87C locus genes (*Hsp70Ba*, *Hsp70Bb*, *Hsp70Bbb*, *Hsp70Bc*) show sustained and progressive induction from 1 to 3 days, identifying them as the import stress-responsive cluster. **(B)** Volcano plot of chaperone genes within the 3 days dataset. The 87C *Hsp70* cluster is massively induced, while small heat shock proteins (*Hsp23*), Hsp90 (*Hsp83*), the CCT/TRiC chaperonin (*CCT2*), J-domain proteins (*DnaJ-60*), the mitochondrial chaperone *Hsp60A*, and *Hsc70-5*/mortalin are repressed, indicating selective remodelling of the chaperone network. **(C)** Volcano plot of mitochondrial ribosomal protein genes within the 3 days dataset. Both mitochondrial large (mt-L) and small (mt-S) subunit genes are broadly downregulated, paralleling the repression of cytosolic ribosomal genes shown in Figure 7E. **(D)** Heatmap of $\log_2$ fold changes for selected genes across 1, 2, and 3 days of mito-IMM$^{DHFR}$ expression. Gene categories include *Hsp70* 87C and 87A loci, chaperones (downregulated), OXPHOS, cytosolic ribosomes, proteasome, antioxidant defence, ISR kinases, eIF2 pathway, V-ATPase, synaptic function, active zone components, and synapse maintenance genes. RNA-seq was performed on dissected larval brains; n = 2 (0 days) or 3 (1–3 days) biological replicates per condition.

**Supplemental Figure 5: PERK overexpression is sufficient to cause synaptic degeneration.**

**(A)**–**(C)** Confocal images of muscle 6/7 NMJs of motoneurons expressing OK371-Gal4 alone (control, **A**), *PEK*-RNAi (**B**), or *PEK* OE (overexpression, **C**). Top row: neuronal membrane (HRP, white) overview. Lower rows at higher magnification: neuronal membrane, Brp (green) and GluRIIc (magenta) merge, and Brp alone. In controls and *PEK*-RNAi expressing animals, Brp and GluRIIc are in precise apposition at all boutons. *PEK* OE causes thinning of NMJ branches, fragmentation of the neuronal membrane, and loss of Brp from individual boutons despite persistent GluRIIc (arrowhead), indicating synaptic degeneration. Asterisks indicate regions shown at higher magnification. Scale bars: 50 µm (overview), 2 µm (zoom-ins). **(D)**, **(E)** Degeneration frequency **(D)** and severity **(E)** at muscle 6/7 NMJs. **(F)**, **(G)** Degeneration frequency **(F)** and severity **(G)** at muscle 4 NMJs. *PEK*-RNAi does not cause degeneration at either muscle type. *PEK* OE significantly increases degeneration frequency and severity at both muscle types. Boxplots indicate median and interquartile range; bars represent mean values with SEM. Data were tested for normality (Shapiro-Wilk) and analysed with parametric or non-parametric tests as appropriate, with post-hoc correction for multiple comparisons. ***$p \leq 0.001$. n = 8 animals with 10 NMJs each (D, F); n = 80 NMJs from 8 animals per genotype (E, G).

**Supplemental Figure 6: GSK2606414 and ISRIB effects at muscle 6/7 NMJs**

**(A)–(B′)** Confocal images of muscle 6/7 NMJs of motoneurons expressing cyt$^{DHFR}$ (**A**, **A′**) or mito-IMM$^{DHFR}$ (**B**, **B′**) (OK371-Gal4), reared on food containing DMSO (mock, **A**, **B**) or GSK2606414 (GSK, **A′**, **B′**). Top row: neuronal membrane (HRP, white) overview. Lower rows at higher magnification: neuronal membrane, Brp (green) and GluRIIc (magenta) merge, and Brp alone. cyt$^{DHFR}$ expressing animals show intact NMJs in both mock **(A)** and GSK **(A′)** conditions. mito-IMM$^{DHFR}$ expression causes synaptic degeneration in mock-treated animals **(B)**. GSK treatment of mito-IMM$^{DHFR}$ expressing animals **(B′)** enhances degeneration. Asterisks indicate regions shown at higher magnification. Scale bars: 20 µm (overview), 2 µm (zoom-ins). **(C)** Schematic of the PERK signalling pathway. **(D)** Frequency of synaptic degeneration. GSK treatment significantly increases degeneration frequency in mito-IMM$^{DHFR}$ expressing animals. **(E)** Severity of degeneration. GSK treatment significantly increases severity in mito-IMM$^{DHFR}$ expressing animals. Boxplots indicate median and interquartile range; bars represent mean values with SEM. Data were tested for normality (Shapiro-Wilk) and analysed with parametric or non-parametric tests as appropriate, with post-hoc correction for multiple comparisons. ns, $p \geq 0.05$, *$p \leq 0.05$, ***$p \leq 0.001$. n = 16 animals with 10 NMJs each per condition (D); n = 160 NMJs from 16 animals per condition (E).

**(F)–(G′)** Confocal images of muscle 6/7 NMJs of motoneurons expressing cyt$^{DHFR}$ (**F**, **F′**) or mito-IMM$^{DHFR}$ (**G**, **G′**) (OK371-Gal4), reared on food containing DMSO (mock, **F**, **G**) or ISRIB (**F′**, **G′**). Top row: neuronal membrane (HRP, white) overview. Lower rows at higher magnification: neuronal membrane, Brp (green) and GluRIIc (magenta) merge, and Brp alone. cyt$^{DHFR}$ expressing animals show intact NMJs in both mock **(F)** and ISRIB **(F′)** conditions. mito-IMM$^{DHFR}$ expression causes synaptic degeneration in mock-treated animals **(G)**. ISRIB treatment of mito-IMM$^{DHFR}$ expressing animals **(G′)** enhances degeneration. Asterisks indicate regions shown at higher magnification. Scale bars: 20 µm (overview), 2 µm (zoom-ins). **(H)** Schematic of the eIF2α signalling pathway. **(I)** Frequency of synaptic degeneration. ISRIB significantly increases degeneration frequency in mito-IMM$^{DHFR}$ expressing animals. **(J)** Severity of degeneration. ISRIB significantly increases severity in mito-IMM$^{DHFR}$ expressing animals. Boxplots indicate median and interquartile range; bars represent mean values with SEM. Data were tested for normality (Shapiro-Wilk) and analysed with parametric or non-parametric tests as appropriate, with post-hoc correction for multiple comparisons. ns, $p \geq 0.05$, *$p \leq 0.05$, **$p \leq 0.01$, ***$p \leq 0.001$. n = 16 animals with 10 NMJs each per condition (I); n = 160 NMJs from 16 animals per condition (J).

#### Supplemental Figure 1

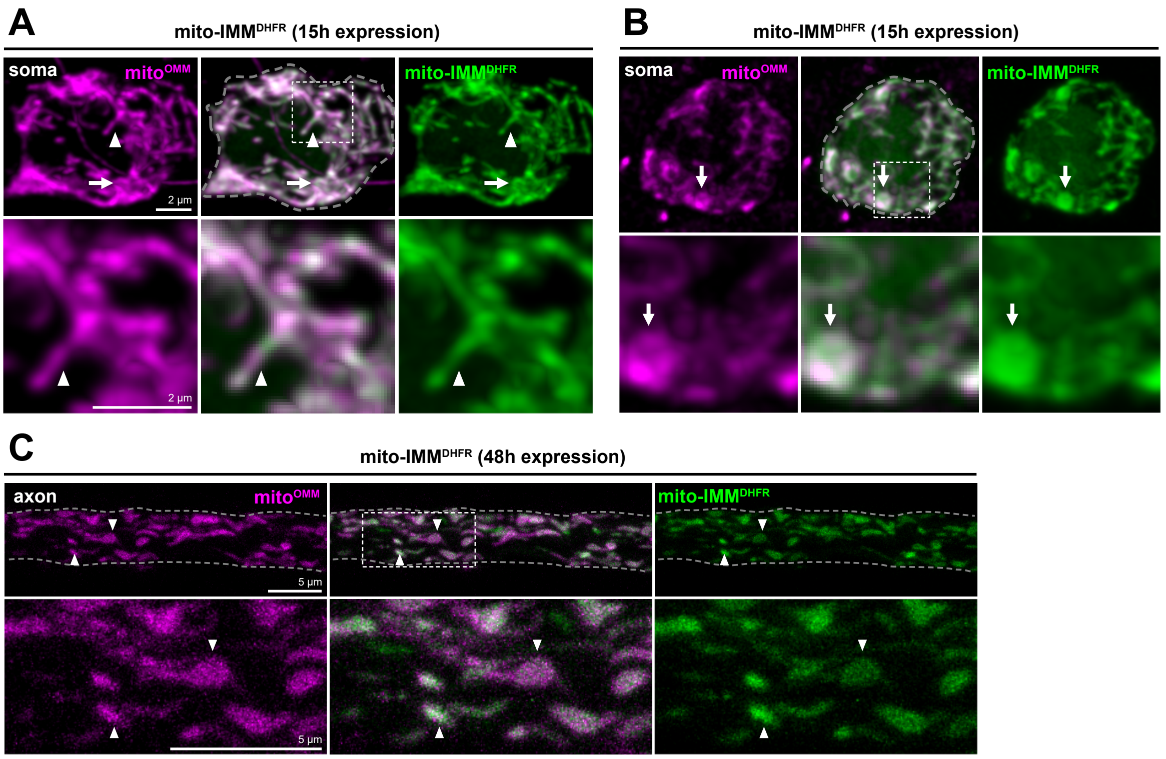

### Supplemental Figure 2

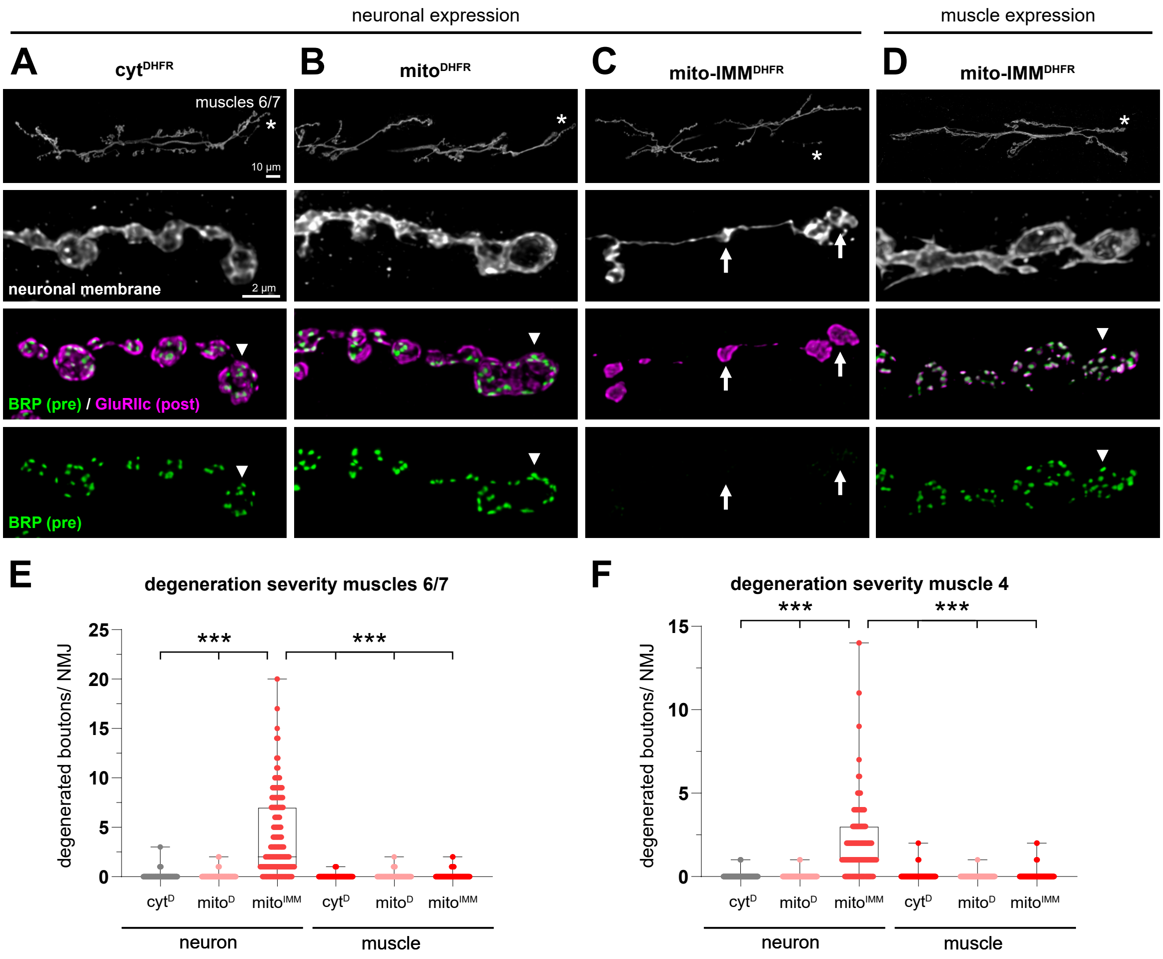

### Supplemental Figure 3

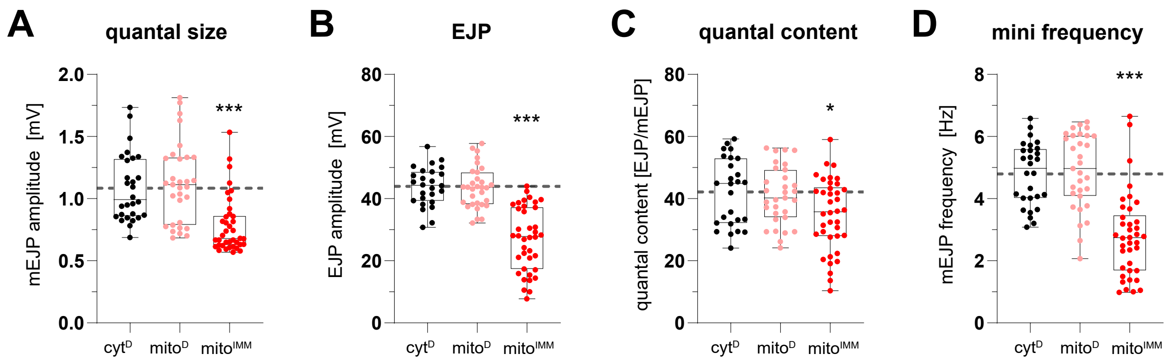

### Supplemental Figure 4

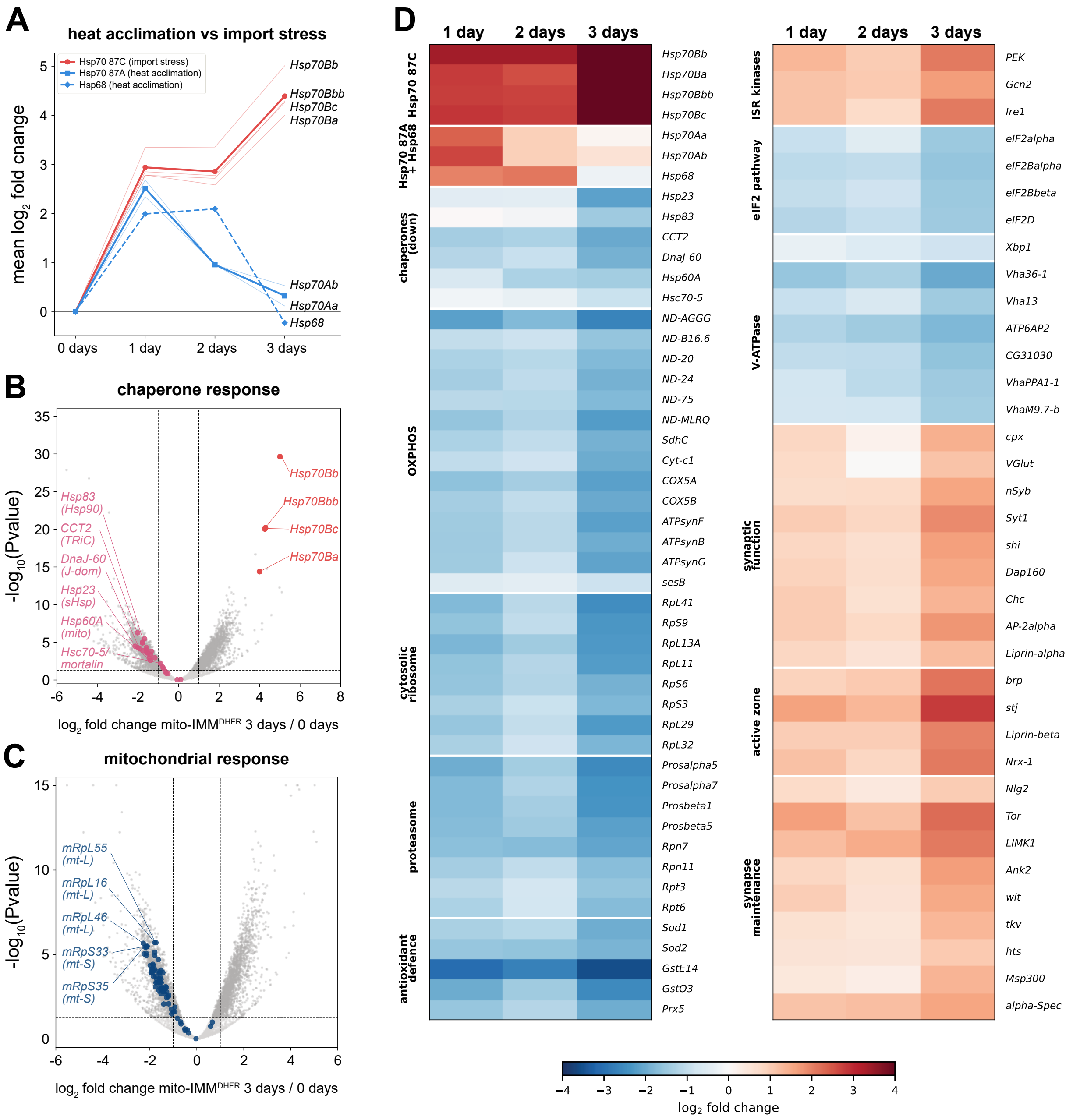

### Supplemental Figure 5

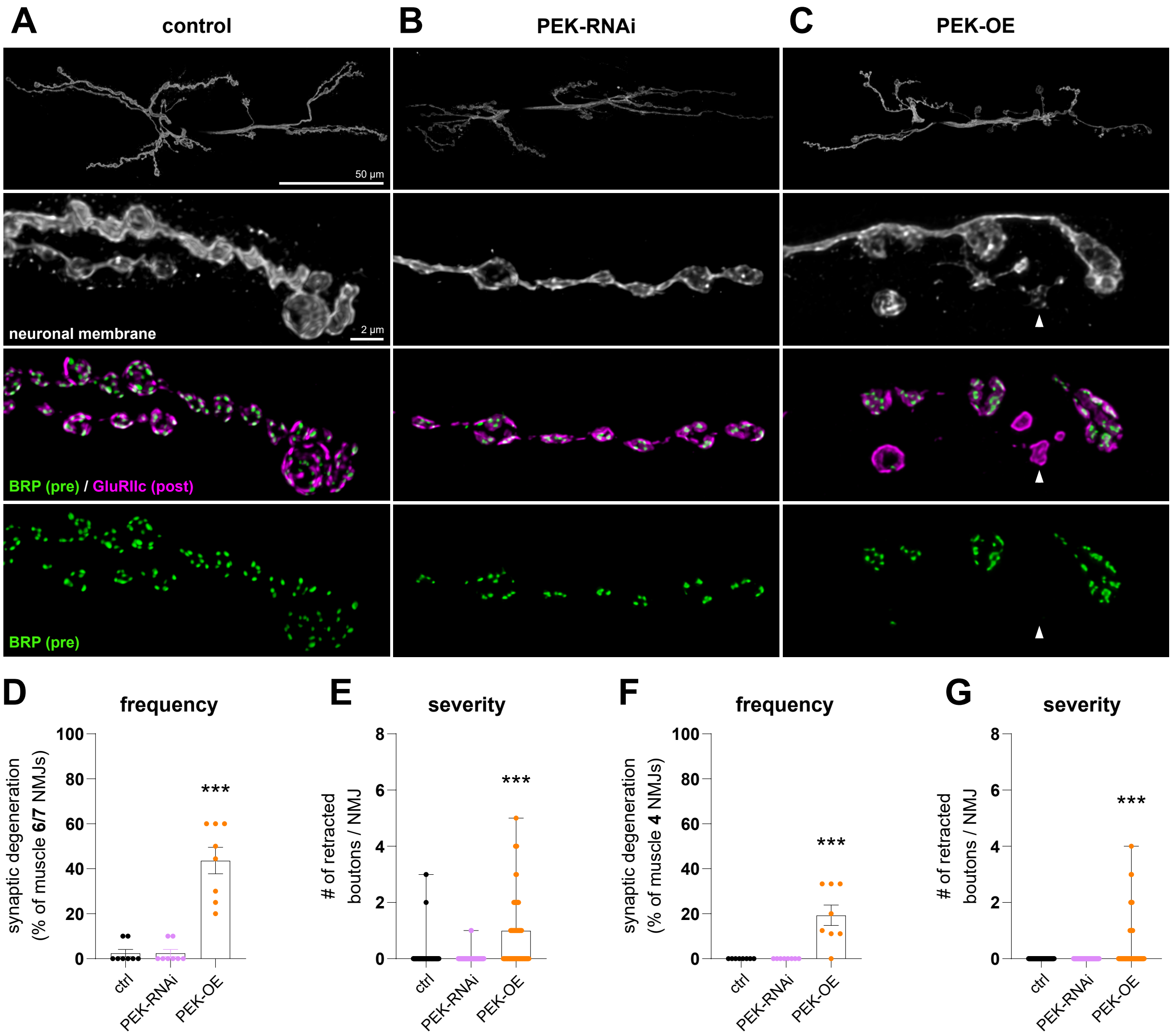

### Supplemental Figure 6

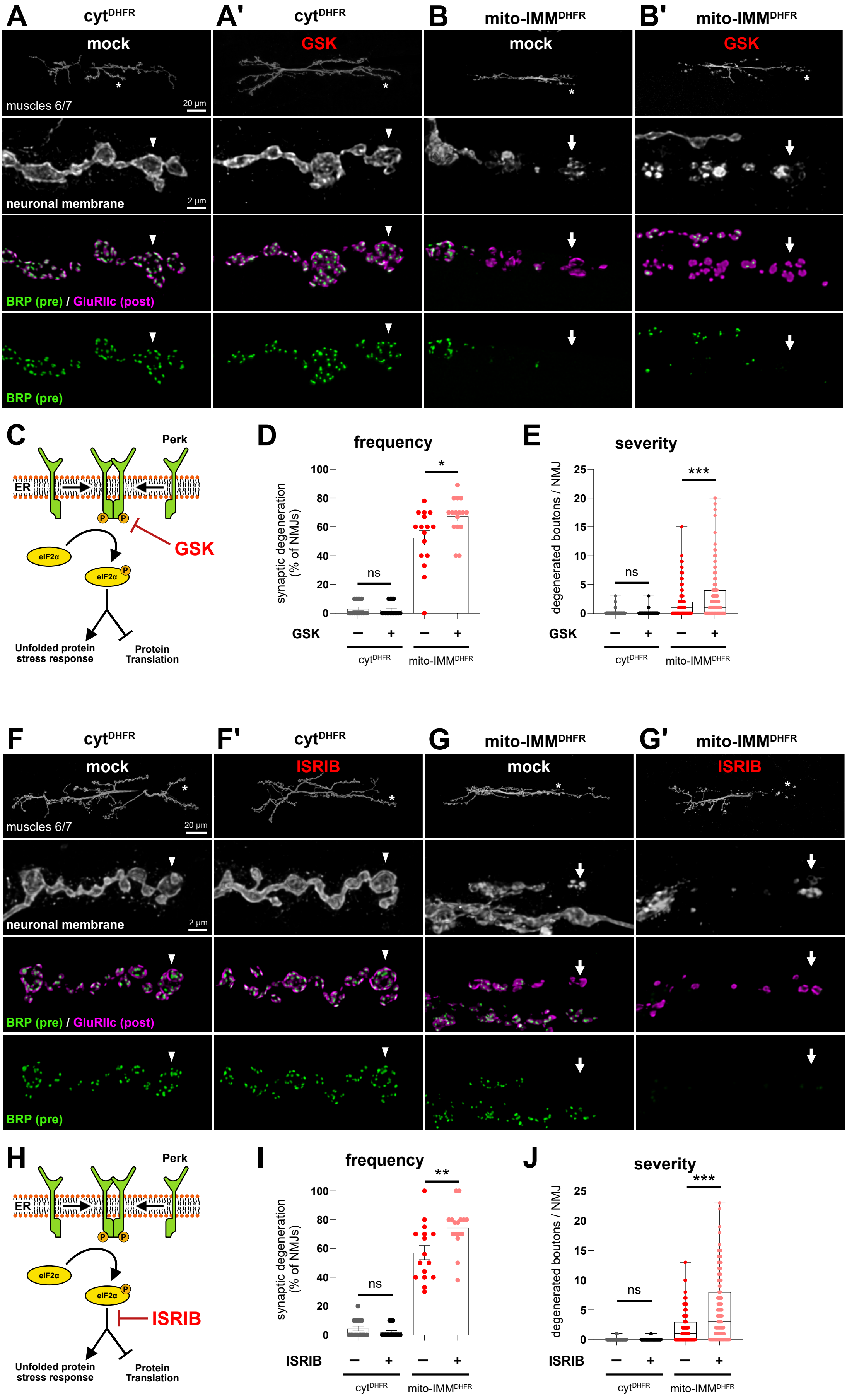
